# Structured sampling of olfactory input by the fly mushroom body

**DOI:** 10.1101/2020.04.17.047167

**Authors:** Zhihao Zheng, Feng Li, Corey Fisher, Iqbal J. Ali, Nadiya Sharifi, Steven Calle-Schuler, Joseph Hsu, Najla Masoodpanah, Lucia Kmecova, Tom Kazimiers, Eric Perlman, Matthew Nichols, Peter H. Li, Viren Jain, Davi D. Bock

## Abstract

Associative memory formation and recall in the adult fruit fly *Drosophila melanogaster* is subserved by the mushroom body (MB). Upon arrival in the MB, sensory information undergoes a profound transformation. Olfactory projection neurons (PNs), the main MB input, exhibit broadly tuned, sustained, and stereotyped responses to odorants; in contrast, their postsynaptic targets in the MB, the Kenyon cells (KCs), are nonstereotyped, narrowly tuned, and only briefly responsive to odorants. Theory and experiment have suggested that this transformation is implemented by random connectivity between KCs and PNs. However, this hypothesis has been challenging to test, given the difficulty of mapping synaptic connections between large numbers of neurons to achieve a unified view of neuronal network structure. Here we used a recent whole-brain electron microscopy (EM) volume of the adult fruit fly to map large numbers of PN-to-KC connections at synaptic resolution. Comparison of the observed connectome to precisely defined null models revealed unexpected network structure, in which a subset of food-responsive PN types converge on individual downstream KCs more frequently than expected. The connectivity bias is consistent with the neurogeometry: axons of the overconvergent PNs tend to arborize near one another in the MB main calyx, making local KC dendrites more likely to receive input from those types. Computational modeling of the observed PN-to-KC network showed that input from the overconvergent PN types is better discriminated than input from other types. These results suggest an ‘associative fovea’ for olfaction, in that the MB is wired to better discriminate more frequently occurring and ethologically relevant combinations of food-related odors.

## Introduction

The cellular basis for associative memory formation and recall remains a central mystery of neurobiology. Connectomics, in which synaptic connections are traced between large numbers of neurons to map circuit wiring diagrams (Lichtman and Sanes, 2008), offers a new method by which to explore the topic. Given the current capabilities of electron microscopy (EM)-based connectomics technologies (Kornfeld and Denk, 2018), the adult fruit fly *Drosophila melanogaster* is arguably an ideal model system for investigating the neuronal networks underpinning learning and memory. Its brain is small enough to have been completely imaged at synaptic resolution by electron microscopy (Zheng et al., 2018); it is behaviorally sophisticated (DasGupta et al., 2014; Dickinson and Muijres, 2016; Ofstad et al., 2011; Owald and Waddell, 2015); and the stereotyped morphology and physiology of its cell types allow ready integration of information across individuals (Costa et al., 2016; Nern et al., 2015). Each cell type normally consists of one or a handful of neurons (Aso et al., 2014; Meinertzhagen, 2010; Scheffer et al., 2020), which may be individually addressed using genetic tools, allowing circuits to be functionally imaged and perturbed in a highly specific fashion (Dana et al., 2016; Dionne et al., 2018; Klapoetke et al., 2014; Venken et al., 2011).

The exception to this norm is the mushroom body (MB; Figure 1A), a bilaterally symmetric structure for associative memory formation and recall (Groschner and Miesenbock, 2019; Guven-Ozkan and Davis, 2014; Heisenberg, 2003). The MB contains about 2,200 intrinsic neurons, called Kenyon cells (KCs), on each side of the fly brain (Aso et al., 2009; Bates et al., 2020; Technau and Heisenberg, 1982). Kenyon cells can be divided into three main subtypes, γ, α’/β’, α/β (Crittenden et al., 1998; Lee et al., 1999; Tanaka et al., 2008), and the axons of each subtype project to the eponymous lobe where the KCs provide input to a relatively small number of MB output neurons (21 cell types comprising 34 neurons, Aso et al., 2014; Aso and Rubin, 2020). Sensory afferents to KCs are dominated by ~150 olfactory projection neurons (PNs), which relay information from the 51 olfactory glomeruli of the antennal lobe (AL; Bates et al., 2020; Jefferis et al., 2007; Stocker et al., 1990; Wong et al., 2002). Projection neuron morphology and odorant response profiles are highly stereotyped across individuals, and exhibit broad tuning and sustained responses to panels of odorants (Bhandawat et al., 2007; Costa et al., 2016). Olfactory PNs project to the rear of the brain and collateralize in the MB main calyx, providing input to KC dendrites. Each KC dendrite terminates in specialized ‘claws’, each of which ensheathes a single PN axonal bouton (Figure 1B). Multiple KC claws commonly ensheath a given PN bouton, and each KC samples input from an average of ~6-8 PNs (Butcher et al., 2012; Caron et al., 2013; Leiss et al., 2009; Yasuyama et al., 2002). Multiple input PNs must be coactive in order to evoke an action potential in a given KC (Gruntman and Turner, 2013), and widefield feedback inhibition is preponderant throughout the MB (Lin et al., 2014), resulting in KC activity that is much sparser and more sharply tuned than PNs (Turner et al., 2008).

**Figure 1.**
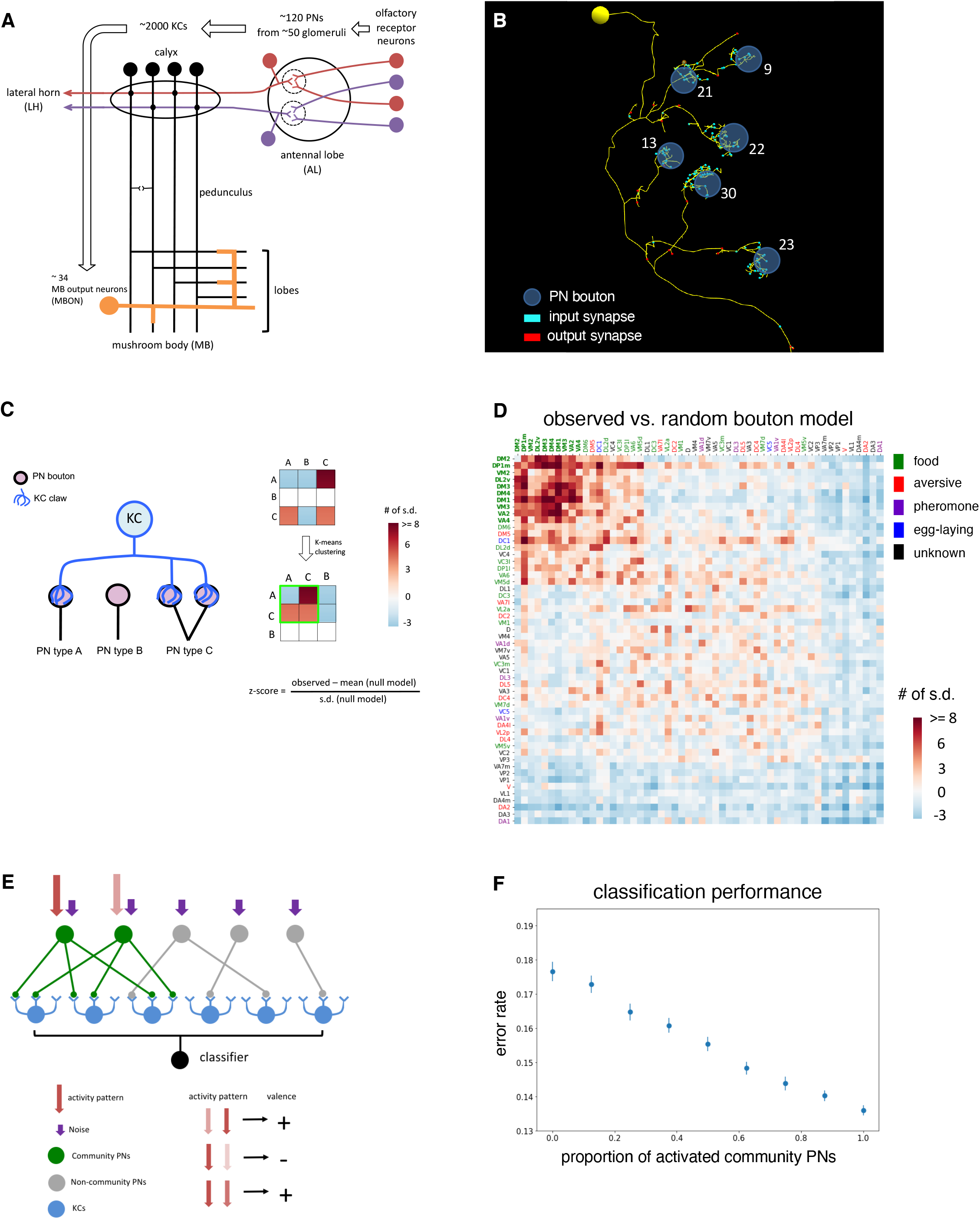
Structured olfactory input to the mushroom body. (A) Schematic of olfactory pathway. Odorants bind to olfactory receptor neurons (ORNs) in the fly antennae and activate a stereotyped subset of glomeruli in the antennal lobe (AL). ORNs in a specific glomerulus provide olfactory inputs to a given class of PNs and the PNs can be classified into ~51 types based on their originating antennal lobe glomeruli. In each hemisphere of a fly brain, ~150 PNs project to two higher brain regions, MB and lateral horn (LH). In the calyx of the MB, the PNs synapse onto ~2,200 KCs. The KCs then converge onto a small number of mushroom body output neurons (MBONS, ~34) at the medial and vertical lobes of the MB. Modification of synapses between KCs and MBONs likely underlies olfactory learning and memory in the fly (Barnstedt et al., 2016; Guven-Ozkan and Davis, 2014; Heisenberg, 2003). (B) Electron microscopy reconstruction of the dendrites of a KC and its olfactory input from PNs in the MB calyx. Kenyon cell dendrites terminate as claw-like elaborations; each claw receives a variable number of synapses (numbers in white) from a single ensheathed PN bouton. Each KC gives rise to a small number of claws (mean ± s.d., 5.2 ± 1.6; Supplemental figure 1). (C) Schematic of conditional input analysis of the PN-to-KC network. Each PN-to-KC connection is treated as binary: if a claw receives three or more synapses from a PN bouton, the KC is considered as receiving one input from that PN type; otherwise it is treated as zero. Input counts across all KCs are then compared to a randomized null model. In the example, given input from PN type ‘A’, KCs are more likely to receive input from PN type ‘C’ and less likely to receive input from PN type ‘B’. A matrix is used to represent the population of these conditional input probabilities. Each row in the matrix represents the probability that, given input from the PN type for that row, KCs are more (red) or less (blue) likely to get input from the PN types in the columns. Re-ordering of the matrix by K-means clustering helps illustrate these relationships. The color scale for each cell in the matrix indicates the z-score, i.e. the number of standard deviations (s.d.) between the observed number of inputs and the mean number of inputs arising from the null model. (D) Structured PN-to-KC connectivity against the random bouton null model. Conditional input analysis was applied to 1,356 randomly sampled KCs on the right side of the fly brain. A specific group of PNs (‘community’ PNs, type names in bold) were found to provide above-chance levels of convergent input to downstream KCs. Olfactory PN types are color-coded according to the category of odorants to which they respond. All community PNs have been reported to primarily encode food-related odors (Supplemental Table 1). (E) Schematic of PN-to-KC network model. In the PN input layer, an olfactory response is represented as a signaling response within a subset of PNs, mixed with Gaussian noise across all PNs. Each of these activity patterns is then assigned a positive or negative valence. Kenyon cell activity is the product of the observed PN-to-KC connectivity matrix and simulated PN activity, subject to a sparseness constraint such that only 5% of KCs are active at any time. The classifier learns to predict the pre-assigned valence based on a readout of KC activity. Performance is quantified by calculating the error rates of the classifier prediction. (F) Using the observed PN-to-KC connectivity, discrimination of inputs from community PNs is superior to that of inputs from non-community PNs. Error bars are standard errors (s.e.) of the mean across 1,000 simulations. For each simulation, a randomly selected subset of PNs are activated. A larger number of activated community PNs leads to better discrimination performance.

The PN-to-KC layer therefore implements a transformation of olfactory representation: broad, stereotyped, and sustained olfactory responses, in a small population of PNs, are converted to sparse, variable, and transient responses, distributed across a large population of KCs. This circuit architecture is an example of a ‘Marr motif’ (Litwin-Kumar et al., 2017; Stevens, 2015), after the theorist David Marr’s foundational work on cerebellar function (Albus, 1971; Marr, 1969). The Marr motif is found in brain regions from different animal species, including cerebellum, hippocampus, and piriform cortex in vertebrates, and even the vertical lobe of the octopus (Cayco-Gajic and Silver, 2019; Farris, 2011; Shomrat et al., 2015; Stevens, 2015). In the fly, it is thought to permit efficient representation of arbitrary combinations of odorants – which may be thought of as points in a high-dimensional olfactory space – for downstream use as a conditioned stimulus during associative memory formation and recall (Cayco-Gajic and Silver, 2019; Groschner and Miesenbock, 2019; Perisse et al., 2013). Theoretical analyses have argued that randomly mixing different input channels, when combined with a nonlinearity such as a spike threshold, increases the dimensionality, and, therefore, the linear separability of activity patterns, making them easier to discriminate (Babadi and Sompolinsky, 2014; Barak et al., 2013; Hansel and van Vreeswijk, 2012; Rigotti et al., 2013). Most models of the PN-to-KC network in the fly have therefore assumed that in the Marr motif, input neurons (PNs) connect to the intrinsic neurons (KCs) at random (Dasgupta et al., 2017; Eichler et al., 2017; Litwin-Kumar et al., 2017; Stevens, 2015; but see Koulakov et al., 2011; Pehlevan et al., 2017).

Several substantial efforts to test the hypothesis of random PN-to-KC connectivity have been made. In the fruit fly larva, a complete PN-to-KC connectome was mapped using a whole-CNS electron microscopy (EM) volume (Eichler et al., 2017). No evidence of network structure was found, although single claw KCs were found to occur more frequently than a gaussian distribution would predict. However, the larval MB is qualitatively and quantitatively different from that of the adult, in that it contains only about 100 KCs per hemisphere (all of which are of a single class γ; Lee et al., 1999). In adult flies, single-cell retrograde labeling was used to identify the PN inputs to a single KC in each of 200 individual flies (Caron et al., 2013). About half the claws for each KC were successfully labeled. No evidence of network structure was found, although some PN types clearly had more downstream targets than others. Finally, electrophysiological recordings of 23 KCs across 27 adult fruit flies revealed highly diverse olfactory responses, with only two KCs exhibiting an identical response profile across individuals (Murthy et al., 2008). Overall, the relatively small sample sizes of the adult datasets have sufficed to exclude highly structured PN-to-KC connectivity graphs, but have not proved randomness.

Indeed, several studies have hinted at the existence of PN-to-KC network structure. Anatomically, PN axonal arbors and KC dendritic arbors are known to occupy stereotyped positions within the MB calyx as a function of cellular subtype (Jefferis et al., 2007; Lin et al., 2007; Tanaka et al., 2004; Zheng et al., 2018). Physiologically, calcium imaging has revealed that KC claws show more correlated responses than would be predicted by chance, and simultaneous optogenetic stimulation of three PN subtypes (comprising ~13 PNs in total) also showed greater-than-chance convergence (Gruntman and Turner, 2013).

Whether the PN-to-KC network is fully random, or has some structure, therefore is an open question. To address it we surveyed a large number of PN-to-KC connections, using the previously described Female Adult Fly Brain (“FAFB”) EM volume (Zheng et al., 2018). The resulting sample of this Marr motif had far greater statistical power than previously obtained datasets, allowing previously undetected network structure to be revealed.

## Results

To map the PN-to-KC network, KCs were randomly selected for reconstruction from a cross-section of the MB pedunculus, a tract where KC axons converge after their dendrites receive input in the MB main calyx (Figure 1A-B; Supplemental Figure 1A-C). The PN bouton innervating each KC claw was then retrogradely traced to the main PN axon trunk, and the PN type was identified, using previously published classifications of PNs in the FAFB dataset (Zheng et al., 2018). Initially, reconstructions were traced purely manually; later reconstructions leveraged an automated segmentation of the full FAFB dataset (Li et al., 2019). All olfactory PN input to 7,102 claws arising from 1,356 KCs was mapped and identified (~62% of all claws on the right side of the brain; 440 KCs were manually traced, and 916 were reconstructed using automated segmentation). Consistent with previous studies (Butcher et al., 2012; Caron et al., 2013), each reconstructed KC was found to have 5.2 claws on average (Supplemental Figure 2A). The number of KCs postsynaptic to each PN subtype was also in excellent agreement with counts obtained from a recently released connectome of adult fly brain connectivity (Supplemental Figure 2B-C; Scheffer et al., 2020). The consistency of these metrics across datasets and methods indicates that the PN-to-KC network reconstructed in the present study is of high quality and therefore suitable for detailed analysis.

If PN-to-KC connectivity were random, the probability that a KC receives input from one PN type is, by definition, independent from whether it gets input from any other type. To test whether these input probabilities are in fact independent, several null models were tested. In the first, the “random bouton” model, a large number of randomized PN-to-KC maps were generated, wherein each claw of each reconstructed KC was assigned a PN bouton selected (with replacement) at random. For each KC, the expected distribution of the number of inputs from each PN type in the random bouton model was obtained. Then, a conditional input analysis was performed, to determine whether KCs are more or less likely than expected to get input from a particular PN type (Figure 1C, matrix columns), given input from another PN type (Figure 1C, matrix rows). Conditional probabilities were quantified as z-scores (the number of standard deviations of the observed value from the mean of the null distribution).

Unsupervised clustering of conditional input probabilities revealed a distinctive ‘community’ of PN types which converge onto KCs at above-chance levels (Figure 1D, PN types in bold). The mean community z score was significantly higher (Supplemental Figure 3A; 5.7 ± 2.9) than non-community PN combinations (Supplemental Figure 3C; −0.5 ± 1.5; Supplemental Figure 3A vs. 3C, p < 1×10^-8^). Additional PN combinations also showed elevated z-scores, but mean z-score for these was significantly lower than the selected subset comprising the community (Supplemental Figure 3B; 2.3 ± 1.9; Supplemental Figure 3A vs. 3B, p < 1×10^-8^). Analysis of individual randomized maps of PN-to-KC connectivity revealed no such clustering (Supplemental Figure 3E). Similar results were obtained using covariance analysis (Supplemental Figure 4).

Following identification of the overconvergent PN community, a literature review was conducted to determine the broad categories of odorants each PN type responds to. Strikingly, all PN types within the community were found to respond preferentially to food-related odorants (Figure 1D; Badel et al., 2016; Hallem and Carlson, 2006; Laissue and Vosshall, 2008; Mansourian and Stensmyr, 2015; Root et al., 2007; Schubert et al., 2014; Semmelhack and Wang, 2009), suggesting that the observed PN-to-KC network structure might play a distinctive role in MB circuit function. Although the MB main calyx contains a great deal of recurrent circuitry (Butcher et al., 2012; Christiansen et al., 2011), with some cell types that are as yet little understood (Zheng et al., 2018), a simplifying feed-forward model of the PN-to-KC network has previously been used to study its performance on classification tasks (Eichler et al., 2017; Litwin-Kumar et al., 2017). When this model was modified to incorporate the observed PN-to- KC network structure, increased activation of community PNs was found to improve classification performance (Figure 1E-F; Supplemental Figure 5). Increased activation of all food-preferring PNs, which includes PN types in addition to the overconvergent community PNs, also led to superior classification performance (Supplemental Figure 5A).

To determine how over-convergence by community PNs is generated, the underlying neuronal network anatomy was further analyzed. Community PN boutons are ensheathed by many more KC claws than expected from the random bouton model (Figure 2A-B). Conversely, fewer KCs than predicted by the random bouton model receive input from the community PN types (Figure 2C). This suggested that the observed network structure might result simply from more ensheathment of community PN boutons by KC claws. To test this hypothesis, a second null model was devised, in which each bouton selects a claw at random (without replacement), and the number of claws ensheathing each bouton is held equal to the observed value. In this “random claw” model, both the number of inputs to each KC and the number of outputs from each PN type are held constant. Clustering of z-scores of the observed PN-to-KC connectivity using the random claw null model revealed the same group of community PNs, albeit with lower variance (Figure 2D; Supplemental Figure 6A-B). Although the random claw model captured more of the observed network structure, over-sampling of the community PNs by KCs (Figure 2A) alone is therefore insufficient to explain the community cluster.

**Figure 2.**
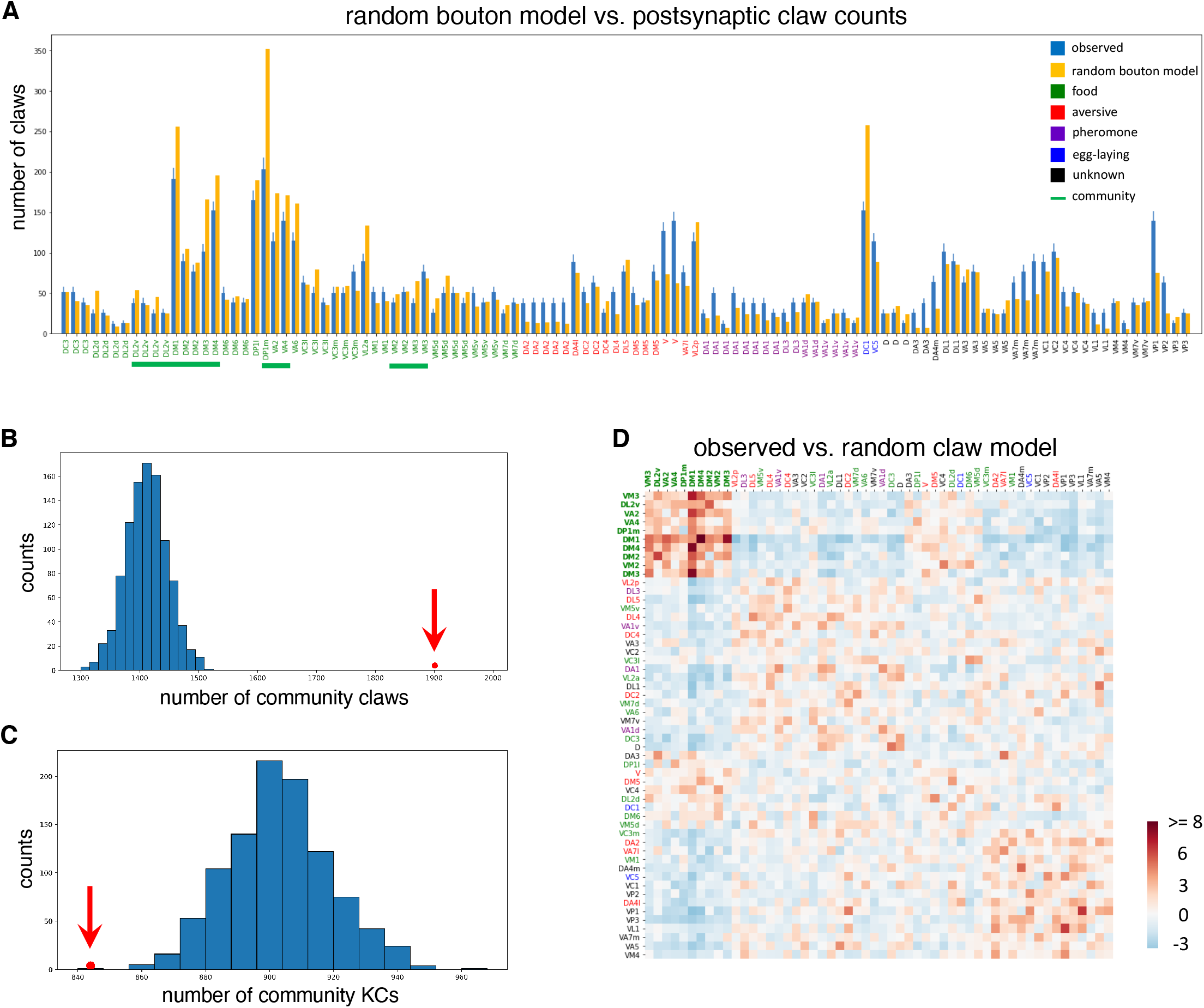
Biased sampling of community PNs by KCs. (A) Biased sampling of PN inputs by KCs. Each bar in the x-axis represents a PN and the y-axis shows the number of claws that receive input from each PN in the random bouton null model (blue) and in the observed network (orange). PNs are grouped by type (i.e. glomerular class) and colored by behavioral significance (as in Figure 1D). Community PN types are underlined. In the observed network, community PNs are usually presynaptic to more KCs than predicted by the random bouton null model (error bars, s.d. of 1,000 random networks; Chi-square test p < 1×10^-10^). (B) Kenyon cells over-sample inputs from community PNs. The observed number of claws receiving input from community PNs (red dot) was greater than the mean of the random bouton null model (distribution of 1,000 random networks; random bouton null model, mean ± s.d., 1412.5 ± 34.0; observed, 1901; z-score, 14.3). (C) Community PNs provide convergent input onto postsynaptic KCs. The observed number of KCs receiving one or more inputs from community PNs (red dot) was lower than the mean of the random bouton null model (distribution from 1,000 random networks; random bouton null model: mean ± s.d., 903.4 ± 17.3; observed, 844; z-score, 3.4). (D) Overconvergent PN-to-KC connectivity contributes to the PN community. Conditional input analysis was applied to the observed PN-to-KC connectivity using the random claw null model, in which PN-to-KC connections are randomized while holding constant both the number of KC claws postsynaptic to each PN type and the number of input PN boutons to each KC. The PN community still emerges, despite the fact that the random claw model incorporates the greater- than-chance output from community PN types (Figure 2A).

In contrast, application of these analysis methods to an earlier sampling of PN-to-KC connectivity (Caron et al., 2013) failed to reveal the community of overconvergent PNs (Supplemental Figure 7A-C). However, that study mapped many fewer PN-to-KC connections (about half the claws in each of 200 KCs; 1 KC mapped per fly). When the data generated in the present study were randomly sub-sampled to match this lower number, minimal network structure was detected and the community could not be discerned (Supplemental Figure 7D). The sample size of the earlier study was therefore likely insufficient to detect the network structure described here.

Both the random bouton and the random claw null models assume that the probability of a PN- to-KC connection is independent of its location in the MB main calyx. However, both PN and KC neuronal arbors are known to occupy stereotyped and circumscribed positions within the calyx as a function of cell type (Jefferis et al., 2007; Lin et al., 2007; Tanaka et al., 2004; Zheng et al., 2018). This suggested that cell type-specific neurogeometry might contribute to the observed nonrandom network structure. Therefore a “local random bouton” null model was constructed, in which each KC claw selects an input at random from the five nearest boutons to it within the MB main calyx (Figure 3A).

**Figure 3.**
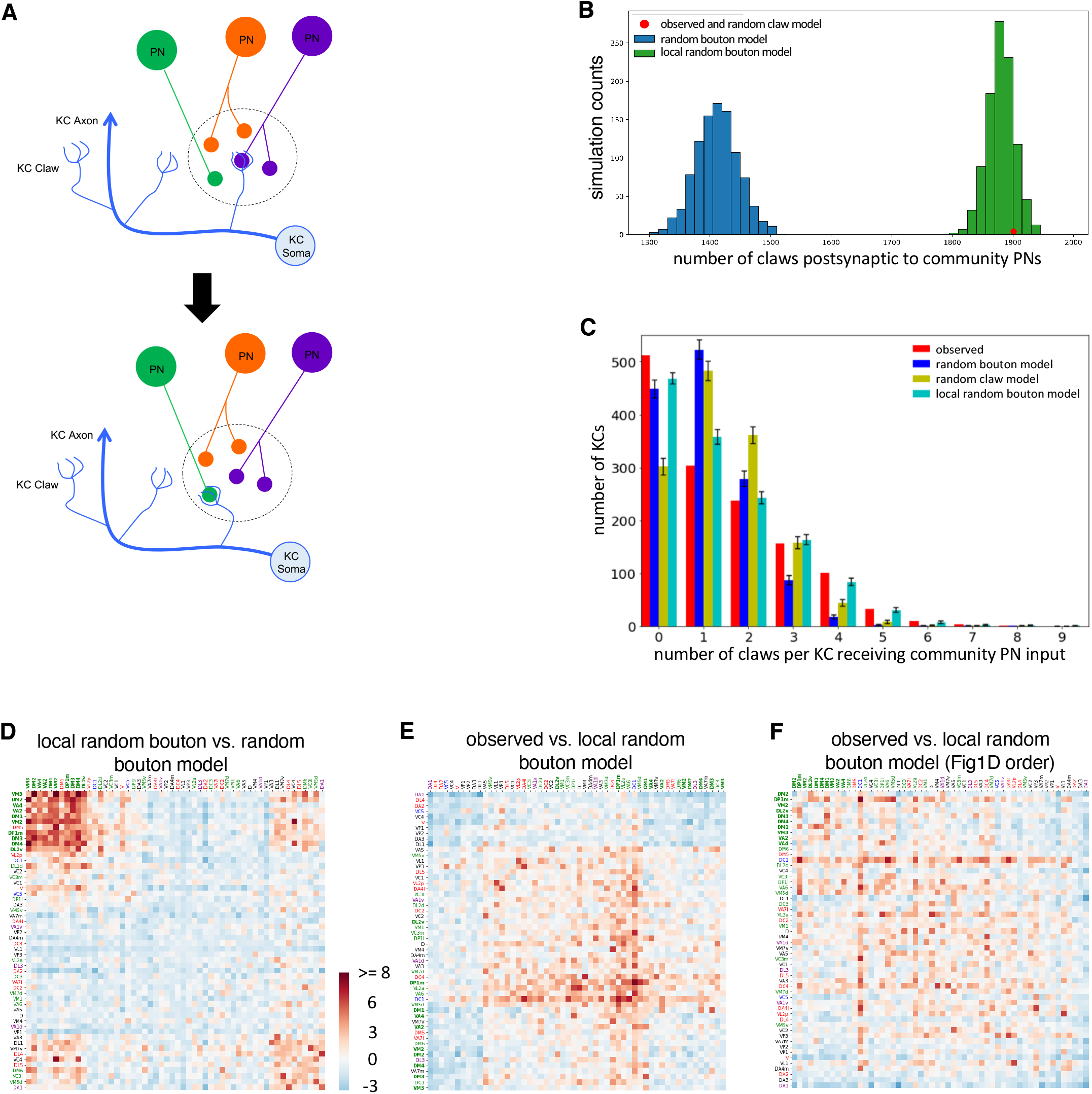
Over-convergence of community PNs to KCs. (A) Schematic for the local random bouton null model. Each claw of each KC is randomly assigned to one of the five nearest PN boutons. In observed PN-to-KC network (upper), one claw from the KC receives input from a PN bouton (purple). The claw is randomly assigned to a different neighboring PN bouton (green). (B) The local random bouton model recapitulates the greater output of community PNs. The observed number of claws receiving inputs from community PNs (red dot) was compared to the number of claws with community inputs in the random bouton model (green histogram) and the local random bouton model (blue histogram). Each distribution represents 1,000 random networks from the null models. Observed (1901) vs. random bouton model (mean ± s.d., 1412.5 ± 34.0), z-scores 14.3. N.b. the random claw null model is constrained to have the observed number of claws postsynaptic to community PNs, and therefore is not considered here. (C) Individual KCs have multiple claws postsynaptic to community PNs boutons. Distributions are shown for the observed network, as well as the means of the random bouton, random claw, and local random bouton null models (error bars, ± s.d.; observed vs. random bouton null model, Chi-square test p < 1×10^-10^; observed vs. random claw null model, Chi-square test p < 1×10^-10^; observed vs. local random bouton null model, Chi-square test p < 0.028). (D) The local random bouton model recapitulates some of the PN community. Conditional input analysis was applied to a representative instance of the local random bouton model, with the random bouton model as the null model. The local random bouton model captures predominantly the same cluster of community PNs (except one PN class, DM5). Community PN types are in bold, and all PN types are color-coded by their response categories as in Figure 1D. (E) Conditional input analysis of the observed connectivity using the local random bouton model as the null model shows no connectivity structure contributed by the community PNs. The remaining network structure is due to biased sampling of other PN types (see Figure 2A). (F) The same matrix shown in (E) but with columns and rows ordered as in Figure 1D. The cluster of community PNs, as seen in Figure 1D, is not seen here, as the community PN network structure is largely recapitulated by the local random bouton null model.

The local random bouton model was superior to the prior models, which lacked spatial constraints. In contrast to the random bouton model, it successfully recapitulated the greater number of claws ensheathing community PN boutons (Figure 3B). It also better recapitulated the overconvergence of community PNs onto KCs. In particular, in the observed PN-to-KC network, some KCs received 3-7 claws of input from community PNs, far more than predicted by chance (Figure 3C, observed vs. random bouton models). Although the random claw model was constrained to preserve the out-degree of each PN type, it was less successful than the local random bouton model in reproducing the observed distribution of multi-claw convergent inputs from the PN community (Figure 3C, random claw vs. local random bouton models). When individual instances of the local random bouton model were compared to the random bouton model, z-score clustering largely recapitulated the observed PN community (Figure 3D); and z-score clustering following comparison of the observed PN-to-KC network to the local random bouton model failed to reveal the PN community (Figure 3E-F).

The success of the local random bouton model suggested that much of the observed non-random network structure arises from the specific neurogeometry of PNs and KCs. Direct visual examination of the community PN axonal arbors and postsynaptic KC dendrites bore out this interpretation. Community PN axons were tightly clustered in peripheral regions of the MB main calyx (Figure 4A-B), and the KCs with the most community input showed dendritic arbors localized to four clusters corresponding to these axonal territories (Figure 4C-E). The four clusters of KC dendrites are consistent with four MB neuroblasts (Ito et al., 1997; Lee et al., 1999). Complete reconstruction of an arbitrarily selected bundle of KCs fasciculating tightly in the MB pedunculus (Supplemental Figure 8) also showed regional bias toward the dorsolateral quadrant of the MB main calyx (Figure 4F), where collaterals of the community PNs tended to ramify. Quantification of pairwise inter-bouton distances revealed that community PN boutons were significantly closer to one another than non-community PNs (Figure 4G). Finally, unsupervised hierarchical clustering divided the PN boutons into 4 distinct territories; one of these clusters was made up of nearly all (9 of 10) of the community PN subtypes (Figure 4F). Thus the community of super-convergent PN subtypes seems to be generated by neurogeometry, as revealed by visual inspection and quantitative analysis of the relevant neuronal arbor structures.

**Figure 4.**
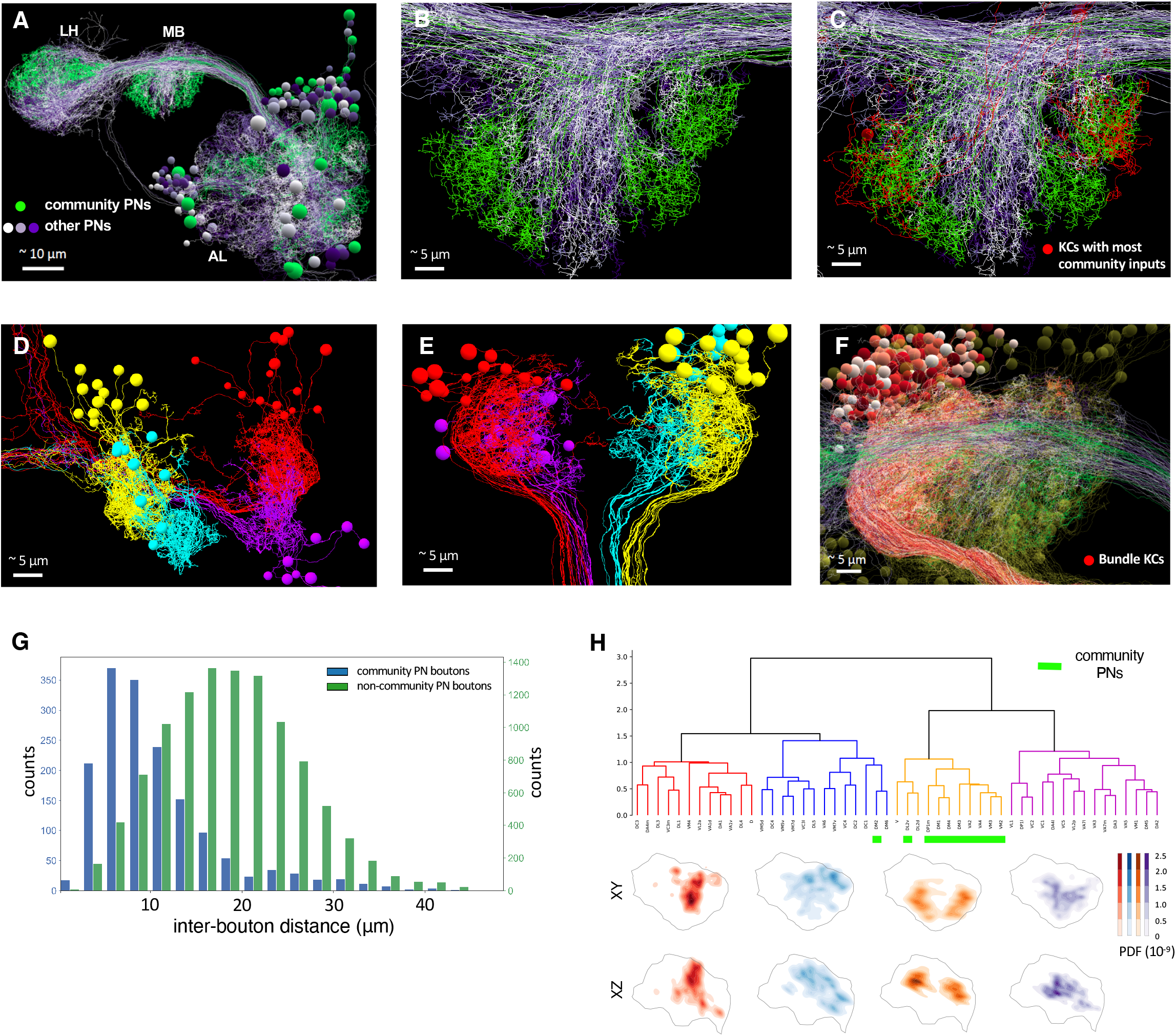
Arbor overlap between subsets of community PNs and KCs. (A) Reconstructed PNs project from AL to two higher brain centers, MB and LH. Community PNs (green) have regionalized projection patterns in MB and LH compared to non-community PNs (white/purple). (B) Frontal view of MB calyx showing reconstructed PN axon arbors; colors as in (A). (C) Same as (B), with the addition of the 6 manually reconstructed KCs receiving 6 or more bouton inputs from community PNs. The dendritic arbors of these KCs (red) overlap with the community PN axon territories (green). (D) Posterior view of MB calyx showing 46 reconstructed KCs that receive 5 or more inputs from community PNs. The dendrites and soma of the KCs, respectively, are segregated into 4 clusters (assigned 4 arbitrary colors) that may correspond to the 4 different neuroblasts of KCs in development (Ito et al., 1997; Lee et al., 1999). (E) Dorsal view of calyx shows 4 different clusters of the same set of KCs as shown and colorized in (D). The cluster axonal bundles also fasciculate in the pedunculus (bottom of figure). (F) Frontal view of calyx shows PN collaterals (colors as in A) and reconstructions of all KCs from a single bundle (“bundle KCs”). The bundle KC dendrites ramify in the dorsal-lateral quarter of the calyx, overlapping extensively with the community PN axonal arbor territory. (G) Community PN boutons are closer to each other than non-community PN boutons. Each count represents the distance between a bouton and its nearest same-type bouton (blue: community PN bouton pairs; green: non-community PN bouton pairs; K-S test p < 1×10^-10^) (H) Unsupervised clustering reveals community PN boutons are spatially colocated in the MB calyx. (upper) Hierarchical clustering based on bouton arbor NBLAST score divides the PNs into four different groups. Nine out of ten community PNs belong to the same group. The y axis of the dendrogram represents Euclidean distances and is cut at 1.5 to divide different PN subtypes into 4 clusters. (lower) Bouton density maps of the four different groups. Colors correspond to the four groups shown in the dendrogram. Color intensity represents density of bouton arbors in a unit space (cubic μm^-1^) and is based on normalized point density function (PDF, Methods).

## Discussion

Our results show that the PN-to-KC network in the adult fruit fly has non-random structure. A community of food-responsive PN subtypes converges at above-chance levels onto downstream KCs (Figure 1D). This network structure is set up anatomically: the axons of participating PN subtypes arborize in restricted regions of the MB main calyx, and the dendrites of many postsynaptic KCs are similarly restricted to those regions (Figure 4). The community PN axonal arbor territories s are similar to those obtained in earlier studies based on light microscopy data (c.f. cluster 1 in Figure 4 C&D Jefferis et al., 2007; Seki et al., 2017; c.f. green cluster in Figure 2 C,E Tanaka et al., 2004). This suggests that the observed PN-to-KC network structure is stereotyped across individuals. The developmental precision required to achieve this structure seems within reach of the fly nervous system, given the highly reproducible geometries of most cell types in the fly brain, including those innervating the MB main calyx (Aso et al., 2014; Lin et al., 2007; Zheng et al., 2018). The PN community we observe in MB is also nearly identical to an independently discovered food-related PN subnetwork formed by axo-axonic synapses between PNs in the lateral horn (c.f. Figure 3F in Bates et al., 2020), suggesting that clustered connectivity of this subset of food-responsive PN types is conserved between brain areas subserving innate (lateral horn) and learned (MB) behavior in the fly.

Why was this structured network connectivity not been seen previously? The likeliest answer may be that past efforts lacked sufficient statistical power to detect the PN community. Subsampling of current dataset to match the number of samples of the most thorough of previous efforts (Caron et al., 2013) renders the community of food-reponsive PNs undetectable (Supplemental Figure 7). Differences in results may also be due to the sampling methods used, but until statistical power is sufficient across all methods, it will be challenging to resolve this question. Furthermore, although our effort is the largest to date, additional network structure may be detected if and when the PN-to-KC network is mapped to completion ipsilaterally and contralaterally in the FAFB dataset. Forthcoming additional brain-spanning EM volumes of the adult fly will also be of interest in this regard (e.g. Scheffer et al., 2020). Alternative analysis methods (e.g. Athreya et al., 2017; Jonas and Kording, 2015; Sporns and Betzel, 2016) might also reveal additional networks structure. It will be of interest to learn whether this community is consistent across individuals, and whether it varies as a function of genetic background, neuronal activity levels, and environmental conditions during development (Kremer et al., 2010; Sugie et al., 2018). Even if the observed network structure is conserved across individuals, it is likely that synaptic output from food-responsive KCs is variable, given that MBON odorant responses are highly variable across individuals (Hige et al., 2015).

What is the functional role of the observed network structure in MB circuit operation? Simplifying models have shown that random connectivity in the PN-to-KC network increases dimensionality and linear separability of neural representation (Litwin-Kumar et al., 2017; Stevens, 2015), indicating that randomly connected Marr motifs may support optimal stimulus classification. However, this assumes that all PNs are activated in a statistically identical fashion. A version of this model incorporating the observed over-convergence of food-responsive PNs onto KCs showed increased discrimination performance for PNs responding to food-related odorants, and decreased performance for the other PN types (Figure 5). This tradeoff calls to mind the efficient coding hypothesis, which states that neuronal resources are allocated to match the distribution of natural stimuli, such that more frequently encountered stimuli are sampled more densely (Barlow, 2012; Laughlin, 1981). In a normal fly’s life, food-related odorant combinations are presumably encountered more frequently than combinations of arbitrary odorants (Mansourian and Stensmyr, 2015). The efficient coding hypothesis predicts that these more frequently encountered combinations of food-related odorants would be sampled more densely than combinations of arbitrary odorants; and indeed, this is what we observe in the *Drosophila* Marr motif. Conceptually, this may be thought of as a kind of ‘associational fovea’, in which more frequently encountered, ethologically relevant combinations of stimuli are sampled more densely (Supplemental Figure 9).

Given the complexity of MB dynamics during learning and recall (Felsenberg et al., 2018; Inada et al., 2017; Owald and Waddell, 2015; Perisse et al., 2016), additional functional characterization of the MB during learning and recall will be needed to determine if the above speculation is correct. Recurrent local microcircuitry is abundant in the MB, including KC-KC synapses (Eichler et al., 2017; Leitch and Laurent, 1996; Liu et al., 2016; Schürmann, 1974), PN-PN synapses (Bates et al., 2020), KC-to-PN synapses (Zheng et al., 2018), and extensive connectivity with local and extrinsic neurons (Amin et al., 2020; Butcher et al., 2012; Christiansen et al., 2011; Inada et al., 2017; Lin et al., 2014; Liu and Davis, 2009). It is also unknown whether the cell types involved fire exclusively in all-or-none fashion, or whether synaptic release can be evoked locally (Zhang et al., 2019). This question becomes especially pertinent given the near ubiquity of mixed input/output neurites in the fly brain (with the exception of the finest dendritic processes) are nearly ubiquitous in the fly brain (Bates et al., 2020; Meinertzhagen, 2018; Olsen and Wilson, 2008; Takemura et al., 2017; our unpublished observations). Downstream of the PN community, different KC subtypes may also play different roles, an aspect not investigated in the present study. Given these complexities, it may be that richer models will be required to fully describe the effect of the observed network structure (Litwin-Kumar and Turaga, 2019).

The present work joins other studies in which unexpected structure is detected in neuronal networks through quantitative comparison of observed connectivity to null models of neurogeometry (e.g. Bopp et al., 2014; Brown and Hestrin, 2009; Egger et al., 2014; Kasthuri et al., 2015; Lee et al., 2016; Mishchenko et al., 2010). Because connectomics data sets offer the exact positions of all synaptic input and output sites on axonal and dendritic arbors, they provide the opportunity to construct unusually well constrained geometric null models. For many classes of neuronal circuit, connectomics data sets may therefore improve the discoverability of network structure compared to alternative methods. This strength among others illustrates how, although connectomics-style wiring diagrams are by themselves clearly insufficient to explain neuronal circuit function (Bargmann and Marder, 2013), they are a useful scaffolding for integrating data across modalities and generating experimentally testable predictions.

## Methods

### Neuron tracing

Neurons were reconstructed from the whole brain EM dataset of an adult fly (Zheng et al., 2018). Skeleton tracing of neuronal arbors and criteria of synapse annotations are conducted as described previously (Zheng et al., 2018) with the CATMAID tracing environment (Schneider-Mizell et al., 2016). To briefly summarize, all the manually traced neurons were reconstructed with an iterative tracing method by at least two tracers, an initial tracer and a subsequent proofreader. The initial tracer reconstructed arbors, followed by systematic review by a different proofreader. When either tracer was not confident about the identifications of a neural process or synapses, they cooperatively examined the image data to reach a consensus. All such sites were further reviewed and resolved by an expert tracer. A chemical synapse was identified if it met at least three of the four following features, with the first as an absolute requirement: 1) an active zone with vesicles; 2) presynaptic specializations such as a ribbon or T-bar with or without a platform; 3) synaptic clefts; and 4) postsynaptic membrane specializations such as postsynaptic densities (PSDs).

Our tracing approach is biased to errors of omission rather than comission. This approach has been shown to have minimal impact on network connectivity in the fly larva (Schneider-Mizell et al., 2016). In addition, the present study is focused on the connectivity between PNs and KCs at a distinctive structure called the microglomerulus, which contains a multitude of synapses between a given PN bouton and its postsynaptic KC claws (Butcher et al., 2012; Leiss et al., 2009; Yasuyama et al., 2002). It is therefore unlikely that the loss of any particular synapse during reconstruction qualitatively affected the analysis described here.

As in Zheng et al. (2018), two reconstruction strategies were used: tracing to classification and tracing to completion. In tracing to classification, in general only backbones and not twigs (microtubule-containing, large diameter neurites, and microtubule-free, fine neurites, respecitvely; Schneider-Mizell et al., 2016) are reconstructed. Tracing is halted once the reconstructed neuronal morphology unambiguously recapitulates that observed by LM or previous EM reconstruction studies for a given cell class. In tracing to completion, all of a given neurite is reconstructed, along with all of its input and output synapses, unless ambiguities in the data make tracing impossible. In some cases tracing to completion is done only within a given brain compartment; in the present study, for example, manually reconstructed KCs were traced to completion only within the MB main calyx (see below).

### Random sampling of KCs

Kenyon cells were randomly sampled from within MB pedunculus ("Random Draw KCs") on the right side of the brain. The pedunculus is a tract of fasciculated KC axons projecting from the posterior of the brain, where KC dendrites ramify in the the MB calyx, to the lobes of the MB at the anterior of the brain, where synapses are made between KCs, MBONs and DANs (Technau and Heisenberg, 1982; Figure 1A). All neuronal processes in a transverse plane of pedunculus (section #4186 in the FAFB dataset) were labelled with seed nodes (2740 in total; Supplemental Figure 1). Seed nodes were randomly selected for reconstruction, which proceeded posteriorly (i.e. retrogradely, in the case of KCs) from the seed node plane. In addition to KCs, the anterior paired lateral (APL) neuron (a wide-field inhibitory neuron; Liu and Davis, 2009), and MB-CP1 (an MBON; Tanaka et al., 2008), were known to have neurites in the pedunculus (Zheng et al., 2018). Therefore tracing to classification was done to determine whether the neuron arising from a given seed node was a KC, using the following morphological criteria. Kenyon cell somata are posterior and slightly dorsal to the MB calyx; each KC makes a handful of dendritic specializations called “claws” within the calyx; and has a single axon projecting anteriorly, with few branches, in the pedunculus (Aso et al., 2014). The APL neuron (one within the MB on each side of the brain) has numerous, densely branching and fine neurites ramifying throughout the entire MB. The MB-CP1 neuron similarly branches densely in the pedunculus and calyx. Disambiguating between these neuron types was therefore relatively straightforward, and tracing was halted if the neuron arising from a seed node was determined not to be a KC. The Random Draw KCs were reconstructed either manually (440 KCs) or by an automatic segmentation-assisted approach (916 KCs), described below. The total sample size of 1,356 KCs was constrained by the time and resources available for the effort; the overall goal was to obtain as large a sample as possible to maximize statistical power.

### Manual tracing of KCs

Each manually reconstructed KC was retrogradely traced to completion from at least section 4186 of the FAFB dataset to the posterior of the brain (some were traced to a greater extent). This spans the posterior ~1/3 of pedunculus and the entire MB calyx. In previous work (Zheng et al., 2018), the boutons of all PNs in calyx as well as the glomerular subtypes of all PNs were identified. Typically, each dendritic claw received input from a single bouton (Leiss et al., 2009; Yasuyama et al., 2002). To facilitate downstream analysis (see below), “claw border” tags were applied to each KC at a node between the “arm” and distal fingers of each KC claw. The “claw border” tags therefore delineated KC claws post-synaptic to distinct PN boutons. Similarly, “bouton border” tags were applied to the PN arbors within MB main calyx.

The majority of reconstructed KCs received olfactory inputs from PNs within MB main calyx. There are 3 main KC classes, γ, α’/β’, α/β, named according to which of the eponymous lobes at the anterior MB the KC axon projects (Aso et al., 2014; Crittenden et al., 1998; Lee et al., 1999; Tanaka et al., 2008). Two additional, numerically fewer types of KC (α/βp and γd) receive non-olfactory inputs such as visual, gustatory, and temperature information via dendritic arbors within MB accessory calyces (Yagi et al., 2016). These were excluded from analysis. All Random Draw KCs were traced to classification anteriorly to section 4186; subtype was assigned depending on which MB lobe the KC axon ramified within.

### Automated segmentation-assisted tracing of KCs

During the KC reconstruction effort, a segmentation of the FAFB dataset became available (Li et al., 2019). A tracing workflow using this segmentation was therefore adopted. Automated segmentation-derived skeleton fragments were manually concatenated, and the entire resulting arbor was proofread as described above. KC claws were only partially reconstructed, sufficient to define which PN bouton was contained and to identify and annotate at least 3 synapses from the bouton to the claw. Control experiments in which one tracing team manually reconstructed KCs to completion and another independently used the automated segmentation to map PN-to-KC connectivity demonstrated the consistency of results between both approaches in quantifying PN bouton/KC claw connection counts (data not shown).

### Conditional input analysis

To determine whether input to KCs from PNs was independent or conditional on PN type, a new method was devised which we termed “conditional input analysis.” The result is a matrix for which a given cell indicates whether, given input from the row PN type, a KC is more or less likely than chance to get input from the column PN type. This approach also allows for detection of asymmetric conditional input (the case where e.g. KCs on average get more input from type C, given input from type A; but less input from type A, given input from type C). Each observed PN bouton-KC claw connection is treated as a single count. The observed number of counts for a given PN type is compared to the distribution of counts generated using a null model. Several null models were used in this study (see below). For each combination of PN types, a z-score is computed (i.e. how many standard deviations from the mean of the null distribution the observed number of counts is). Unsupervised K-means clustering of the z-score matrix was used to group matrix entries.

A summary of the steps in conditional input analysis follows; source code is available at https://github.com/bocklab/pn_kc.

Projection neuron types are named after the glomerulus (‘Glom’) in the antennal lobe that PN’s dendrites innervate. Consider types Glom A, B, C, and so on. For a given connectivity matrix,

1. Select all KCs having at least one claw receiving input from a bouton of Glom A.
2. The number of inputs to these KCs from Glom B, C, D, and so on are counted. This provides a count of the number of inputs to the KC cell population from Glom B-D, given input from Glom A.
3. Repeat (1) – (2) for Glom B, C, D, and so on.
4. For each null model (see below), repeat (1)-(3) above on 1,000 *in silico* randomizations of the observed PN-to-KC network. This generates the null distributions from which a z-score can be generated for observed connectivity for each PN type pair. A matrix of these z-scores is termed a “conditional input matrix”.
7. Apply K-means clustering to the conditional input matrix. The K-means algorithm (MacQueen, 1967) clustered PN types into groups with equal variances and the cluster number of each PN type is used to re-order both the columns and rows of the z-score matrix.

K-means clustering of the conditional input matrix groups glomeruli with similar z-scores together, and therefore reveals subsets of PNs that provide more (or less) input than predicted by a given null model. Over-convergence of inputs (red in our figures) is more strongly detected by this approach, since the random bouton null model (see below) can result in PN types having a small number of boutons to have zero KC outputs. This, in turn, lowers the magnitude of negative z-scores (since the mean of the null model values is already low).

### Null models of PN-to-KC connectivity

Three null models were used: (1) random bouton model, (2) random claw model, and (3) local random bouton model.

In the *random bouton model*, each Random Draw KC claw is reassigned, with replacement, to a randomly selected PN bouton in the calyx. On average, therefore, the number of outputs provided by PN type (i.e. out-degree per PN type) will be proportional to the number of boutons that belong to that type. The number of claws for each KC (i.e. in-degree per KC) is also maintained. To apply conditional input analysis to the data of Caron et al. (2013) using this null model, the bouton counts per PN type obtained from the present work were used (Supplemental Figure 7), since bouton counts per PN type were not generated in that study.

In the *random claw model*, each PN bouton is reassigned claws at random, without replacement. The number of claws so assigned is equal to the number of claws ensheathing that bouton in the observed PN-to-KC network. Thus in this randomization, the number of claws receiving input from a given PN type (i.e. out-degree per PN type) and the number of claws each KC has (i.e. in-degree per KC) are maintained.

In the *local random bouton model*, each claw of each KC is randomly assigned to one of its five nearest boutons (including the one it ensheathed in the observed network), with replacement. Distances were measured between claw and bouton centroids. In this randomization, KC in-degree and geometric constraints on connectivity are preserved.

### Covariance analysis of connectivity

Covariance analysis (Newman, 2018) is a commonly used measure of whether two inputs occur more frequently than predicted by chance and as such is an alternative to the conditional input analysis described above. Its output is a matrix of p-values of input rates compared to the expected distribution arising from given null model. The procedure is summarized as follows.
1. The covariance measure for each pair-wise combination of PN types was computed for the observed connectivity.
2. The observed PN-to-KC connectivity was randomized 1,000 times using the random bouton model. For each randomization, a covariance matrix of PN types was computed.
3. For each pairwise combination of PN types, a p value is estimated by counting how often the randomized covariance was great than or equal to the observed covariance. A p value of less than 0.05 (significance level) implies the probability of obtaining such a covariance in a random network is low, and the alternative hypothesis of seeing such an observed value in a null model is therefore rejected. The results are shown in a p-value matrix (Supplemental figure 4 A - D) in which each cell represents a p value for a given pair of glomeruli indicated in the corresponding row and column labels.
4. The p-value matrix was re-ordered either using the Fig 1D clustering order (Supplemental Figure 4 A, C) or using order given by K-means clustering (Supplemental Figure 4 B, D). To cluster statistically significant but numerically small p values, K-means clustering was performed on a binary version of the p-value matrix wherein all p values less than 0.05 were set to 1, and otherwise to 0.

For the analysis of synaptic connectivity (Supplemental Figure 4C, D), covariance measures were directly calculated from synapse counts, using only the manually reconstructed Random Draw KCs (whose dendritic arbors in MB calyx were reconstructed to completion; see Manual Tracing of KCs, above). To generate the null model of synaptic connectivity, the bouton-claw binary network is randomized and each bouton-claw connection is assigned a synapse count that was randomly drawn (with replacement) from the distribution of number of synapses per claw.

### Clustering analysis of PN boutons

Each PN type was classified and each the bouton in MB calyx was annotated in previous work (Zheng et al., 2018). Using these annotations, skeleton reconstructions of each bouton were extracted. Pairwise NBLAST scores on the bouton skeletons were computed (Costa et al., 2016) and clustered by Ward’s algorithm (Murtagh and Legendre, 2014). NBLAST is a similarity measure for both shape and position; in this case, because the skeletons within each bouton were small, clustering is likely based mostly on bouton position. The probability of bouton arbors being in a given location in calyx was estimated following the approach of Bates et al. (2020). In brief, the bouton skeletons were resampled evenly at 0.1 μm intervals. A Gaussian kernel density estimate (KDE) was used to fit the number of skeleton nodes per unit space (cubic μm^-1^) for each of the two projected dimensions (x,y or x,z). The density map therefore reflects the probability (point density function, PDF) for boutons of a given PN type to be found at a given location in MB calyx. The PDF is normalized to the same scale (0 – 2.5 × 10^-9^) for each of the four groups. The boundaries of MB calyx were generated from an nc82 (synapse)-stained template brain aligned to the FAFB image volume as described in Zheng et al. (2018).

### Comparison of postsynaptic KC counts between FAFB and hemibrain datasets

During preparation of this manuscript, a segmentation of a portion of a second adult fly brain became available in preprint form (the ‘hemibrain’; Scheffer et al., 2020). In the hemibrain dataset, all PNs and ~2,000 KCs on the right side of the brain were segmented as part of a large-scale proofreading effort (50 person-years over ~2 calendar years). As of this writing, the publicly available hemibrain segmentation does not demarcate PN bouton and KC claw boundaries, preventing straightforward application of our analysis approach. We used the hemibrainr package (https://github.com/flyconnectome/hemibrainr) to download the connectivity matrix between all PNs and KCs from the dataset server (hemibrain v. 1.0.1, https://neuprint.janelia.org). The connectivity matrix is then binarized such that each unique pair of PN and KC with 3 or more synapses is defined as one connection and otherwise zero. For a PN, the number of connections is equivalent to the number of KCs postsynaptic to the PN. Connections for different PNs of a common type are summed and divided by the total number of connections in the entire binary connectivity matrix. The percentage of connections for each PN type is used to compare to the same number from FAFB (Supplemental Figure 2B-C).

### Modeling

The PN-to-KC network model was a modification of earlier models used in the larval and adult fly (Eichler et al., 2017; Litwin-Kumar et al., 2017, respectively). In these models, simulated activities across all PNs are created for each stimulus odor. Each stimulus is randomly associated with one of two categories with equal probability. The PN activity (signal) is generated by drawing independently from a rectified unit Gaussian distribution corrupted by Gaussian noise (s.d. 0.2). To probe the effect of the observed overconvergence of community PN types (10 types comprising 16 individual PNs), for each stimulus,16 modeled PNs were activated (i.e. Gaussian activity patterns were created). Within the 16 activated PNs, the fraction of community PNs was varied as follows: 0 (16 non-community PNs), 0.125 (2 community PNs, 14 non-community PNs), 0.25 (4 community PNs, 12 non-community PNs), 0.375 (6 community PNs, 10 non-community PNs), 0.5 (8 community PNs, 8 non-community PNs), 0.625 (10 community PNs, 6 non-community PNs), 0.75 (12 community PNs, 4 non-community PNs), 0.875 (14 community PNs, 2 non-community PNs), 1 (all 16 community PNs). These fractional values comprise the x-axis of Figure 1F. For fractional values less than 1, activated PNs were randomly selected from the 16 community PNs. For all fractional values, non-community PNs were randomly selected from the population of 97 non-community PNs. Gaussian noise with standard deviation 0.2 was then added to the activity levels of all PNs. Kenyon cell activity is given by multiplying PN activity by the matrix of PN-to-KC connections *m* = *Θ* (*h* – *θ*), where *θ* is a threshold whose values are picked for each KC such that each KC is active with a probably of *f*(f = 0.05, also called coding level) for each stimulus. The *Θ* is a rectification term. The *h* represents input activity provided by each PNs multiplied by their corresponding number of synapses to the KC. KC activity patterns were used to train a maximum-margin classifier, and the goal is to predict the pre-assigned one of two categories. In the testing phase, the same set of PN activity patterns corrupted with different applications of Gaussian noise were used as input, and the resulting KC activity patterns were given to the trained classifier to predict which of the two categories each stimulus belongs to. Error rates of the prediction from 1,000 simulations were used to evaluate classifier performance. Because this is a two-alternative classification task, the expected error rate for chance performance is 50%. For reporting error rate results (Figure 1F, Supplemental Figure 5A-B), standard errors of the mean (s.e.) are used as the goal is to compare mean error rate of different models.

The model requires synapse counts between each connected PN-KC cell pair. In Figure 1F and Supplemental Figure 5A, the synaptic counts between PNs and manually reconstructed Random Draw KCs were used. In Supplemental Figure 5A, the same model is implemented except that 38 food PNs (Supplemental Table 1) are chosen to be activated. In the activated food-PN datum (red dot), 38 food PNs are activated with simulated Gaussian activity patterns and all PNs, including the food PNs, are corrupted with Gaussian noises (s.d. 0.2). In the null model distribution (blue histogram in Supplemental Figure 5A), for each simulation (one count in the histogram), a random set of 38 PNs are picked to be activated and error rates of the classifier are computed to evaluate performance of the models. In Supplemental Figure 5B, each KC’s connections to a given PN were randomly reassigned (with replacement) to different PN, and 16 PNs, with varying proportion of community PNs, are activated using the same model implementation as in Figure 1F.

### Statistics

When comparing two or more distributions, if the data are categorical (e.g. Figure 2A, 3C, Supplemental Figure 2C) a Chi-square test is used. When the data are continuous (e.g. Figure 4G, Supplemental Figure 3B-D, 6B), a Kolmogorov–Smirnov test (K-S test) is used. When a distribution is compared with a observed datum (i.e. a single data point), as in each cell of the conditional input matrices (e.g. Figure 1D, 2D, 3D-F, Supplemental Figure 3E, 6, 7A, C-D), in Figure 2B, C, 3B, and in Supplemental Figure 5A, a z-score (see Conditional input analysis, above) is computed.

---

**Supplemental Figure 1.**
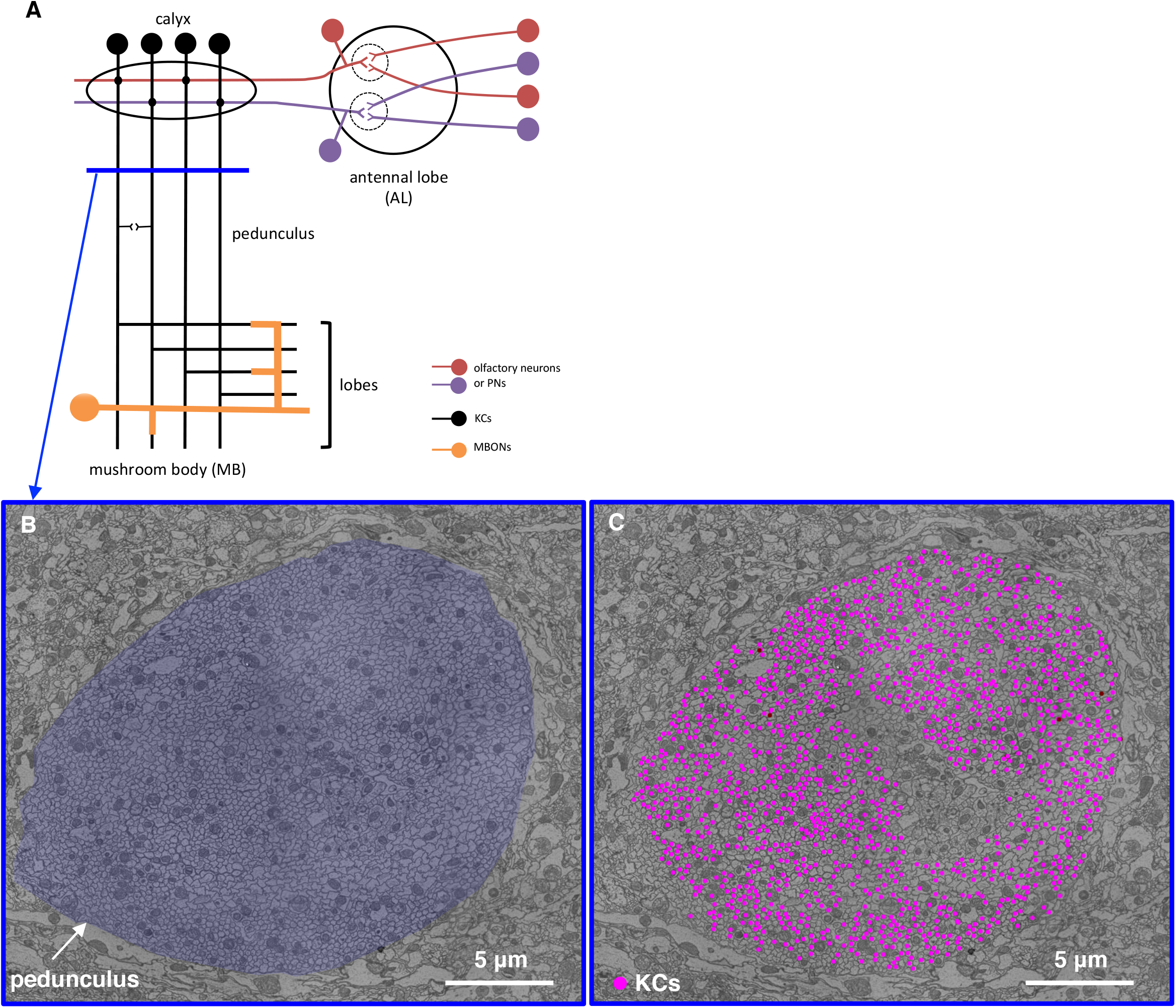
(A) Schematic of MB anatomy, as in Figure 1A. Kenyon cell axons fasciculate and project anteriorly in parallel within the pedunculus. The blue line in the pedunculus indicates the location of the transverse plane where KCs were randomly sampled for reconstruction (B - C). (B) Subarea of a frontal section from the whole-brain EM volume, showing the cross-section through pedunculus (blue false color) used for random sampling (C). (C) Randomly sampled KCs in the pedunculus. The cross-section of each randomly sampled KC axon is annotated with a magenta dot. All neurite cross-sections within the pedunculus were initially annotated (not shown); if a neurite was randomly sampled for reconstruction that turned out not to be a KC, it was discarded from further analysis. N.b. a discrete region in the middle of the pedunculus is occupied by other cell classes such as APL and non-olfactory KCs from accessory calyces (i.e. KC-α/βp and KC-γd), hence there are no magenta points in this region.

**Supplemental Figure 2.**
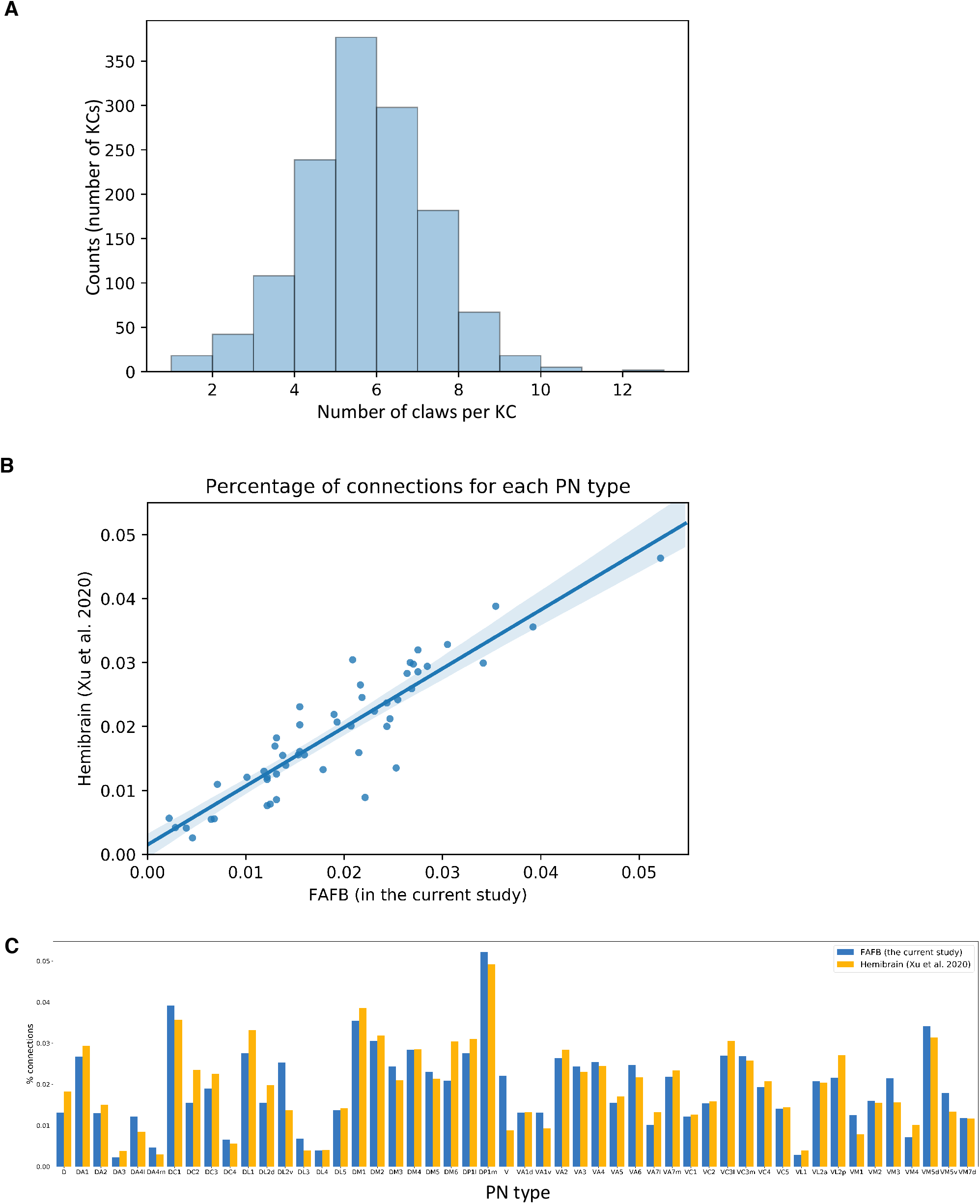
(A) Distribution of number of claws per KC for all randomly sampled KCs (mean ± s.d., 5.2 ± 1.6). (B) The number of postsynaptic KCs per PN type is consistent between the current study and the connectome deriving from the recent ‘Hemibrain’ dataset (v. 1.0.1; Scheffer et al., 2020). Each point represents a PN type; three or more synapses between a unique PN-KC pair is counted as an individual PN-KC connection. Since the two datasets have different numbers of reconstructed KCs, output from each PN type is represented as a percentage. There is a tight correlation across the two datasets (*r*^2=^0.83; blue-gray, 95% confidence interval along the regression line). (C) The same data as in (B), with PN types identified.

**Supplemental Figure 3.**
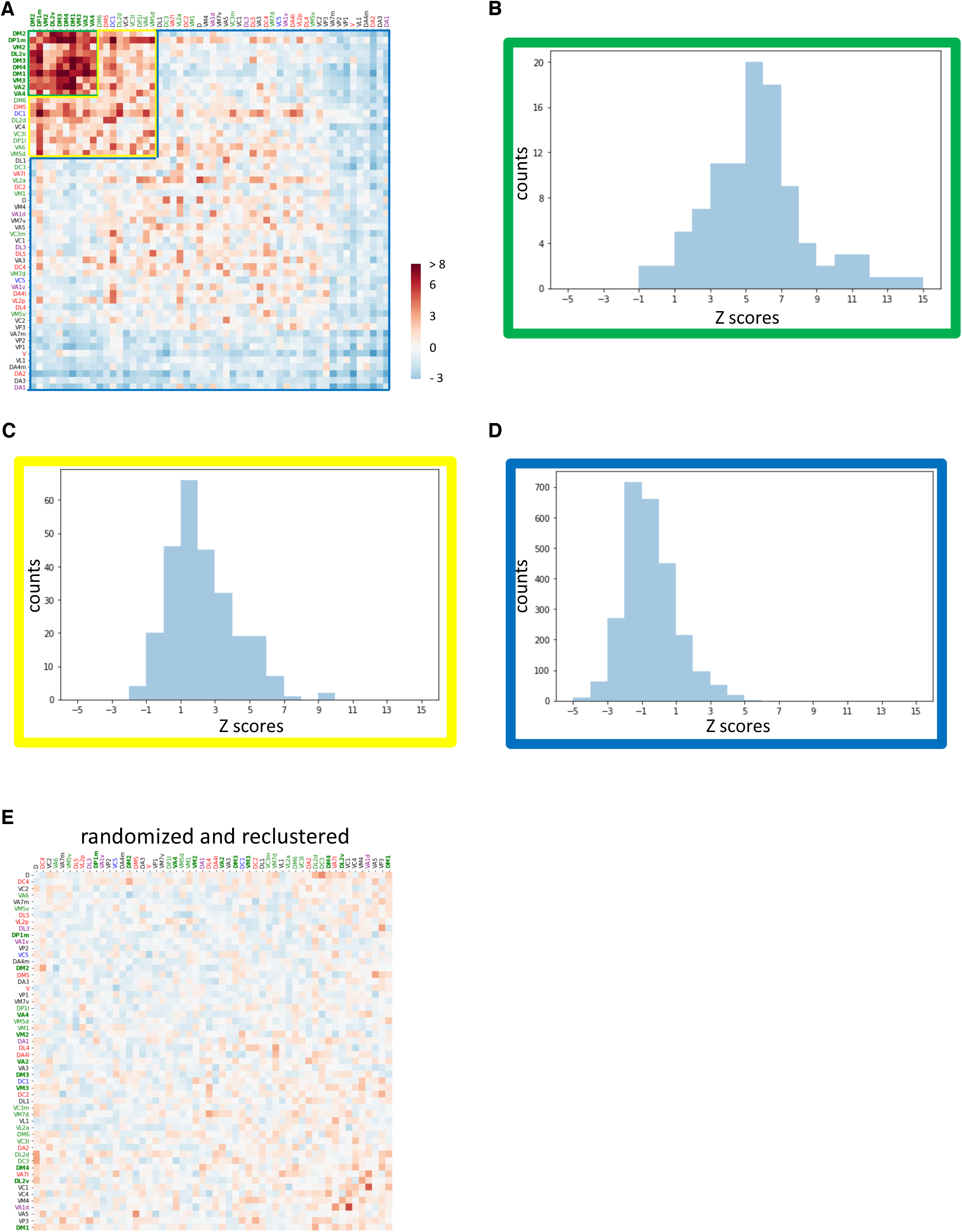
(A) Figure 1D, with colored boundaries delineating the matrix subregions for which z-score distributions are shown in (B-D). The distributions are significantly different (B, green; C, yellow; D, blue). (B) Distribution of z-scores for community PN types (green area in panel A). (C) Distribution of z-scores for PN types weakly clustering with the community PNs (yellow area in panel A). It is significantly different from the community PN distribution (K-S test p < 1×10^-10^). (D) Distribution of z-scores for remaining PN types (blue area in panel A). It is significantly different from the community PN distribution (B, above; K-S test p < 1×10^-10^) and the weakly clustering PN types (C, above; K-S test p < 1×10^-10^). (E) Conditional input analysis of a single representative network from the random bouton model, shows no clustered structure in the z-score matrix. The random bouton model was also used as the null model. Any connectivity structure that deviates from the null model will manifest as clusters of high or low z-scores (2 s.d. or more as compared to the mean of the null model) in the matrix. No discernible cluster is seen after re-clustering of the z-score matrix, showing that the observed clustering is unlikely to be an artifactual result from an expected distribution of random values.

**Supplemental Figure 4.**
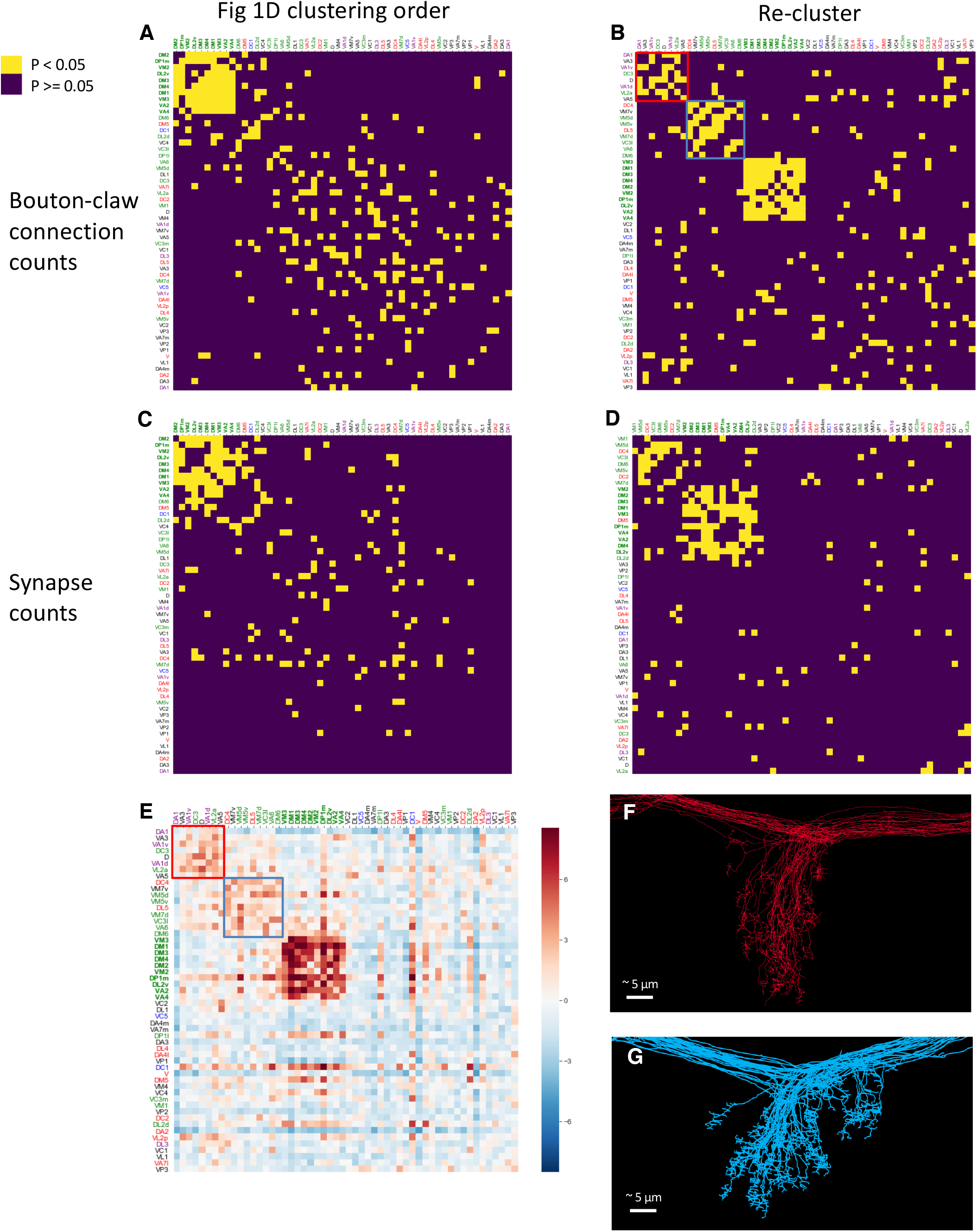
(A) Co-variance analysis (Methods) of the observed PN-to-KC connectivity. This approach generates a symmetric matrix of p-values for PN type combinations. The lower the p-value, the less likely the observed convergence of the PN type pair onto postsynaptic KCs is expected to occur by chance. Values < 0.05 are color coded in yellow; others are black. The column-row ordering of PN types is the same as in Figure 1D. The PN community is discernible as a mostly yellow square at the top left. PN type response categories are color coded as in Figure 1D. (B) As in (A), except the covariance matrix is reordered using unsupervised K-means clustering on p-values. The PN community (type names in bold) is reconstituted following this reclustering. Two weaker clusters of PN types (red and blue squares, overlaid) are discernible. (C) As in (A), except covariance between synapse counts between each PN and KC pair was quantified. The PN community is still discernible. (D) As in (B), except covariance between synapse counts between each PN and KC pair was quantified. As with (A-C), the PN community types comprise the dominant cluster, confirming that the main finding is robust to different analysis methods. (E) The z-score matrix from the conditional input analysis (Figure 1D) is re-ordered with the order given by K-means clustering of the co-variance matrix as shown in (B). The re-ordering reveals the same two weaker clusters of PNs as seen in (B). (F) Anterior view of MB calyx shows axon collaterals of reconstructed PNs of the types shown in the top left weak cluster in (B), demarcated by a red square. The PN collaterals occupy a conscribed territory within the MB main calyx. (G) As in (F), except the PN types are from the second weak cluster in (B), demarcated by a blue square.

**Supplemental Figure 5.**
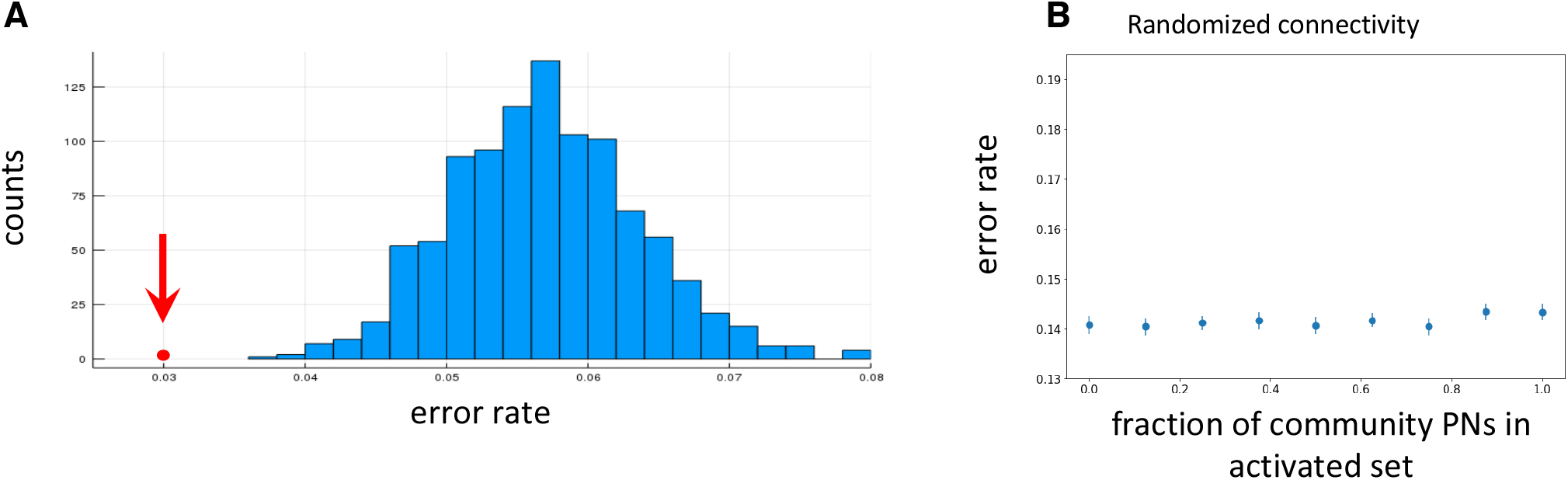
(A) In a model using the observed PN-to-KC network, stimulus discrimination by food odorant-responsive PNs is superior to discrimination by other PN types. The classifier seeks to discriminate different sets of PN activity patterns based on the KC responses, which are integrated via the observed PN-to-KC connectivity (for schematic see Figure 1D). In each set of PN activity patterns, 38 PNs are activated. The red dot (x=0.03) indicates average performance from 1,000 simulations of the model with activation of all food-responsive PNs (38 in total, including PN types that are not part of the community identified in Figure 1D). The distribution (mean 0.057, s.e. 0.007) shows error rates from 1,000 sets of 38 non-food PNs that are activated. Each data point represents the average error rate for a set of 38 PNs randomly selected from all non-food PNs. The number of non-food PNs that are activated (38) is kept consistent with the total number of food PNs. Observed vs. blue histogram, z-score - 4.0, p < 1×10^-4^. (B) With a randomized PN-to-KC network, activating varying fractions of community PNs results in unchanging classification performance. Each KC claw is randomly assigned to a PN with equal probability for each PN, and each PN-claw pair is assigned a number of synapses randomly drawn from the distribution of synaptic counts for all manually reconstructed claws. For each plotted data point, the same number of PNs (16) is activated, but with a varying fractions of community PNs (indicated in x-axis). Each data point is the average of 1,000 simulations, each of which represents a combination of randomly selected community PNs and non-community PNs according to the indicated fractions (e.g. in the case of 0.5, 8 randomly selected community PNs and 8 randomly selected non-community PNs).

**Supplemental Figure 6.**
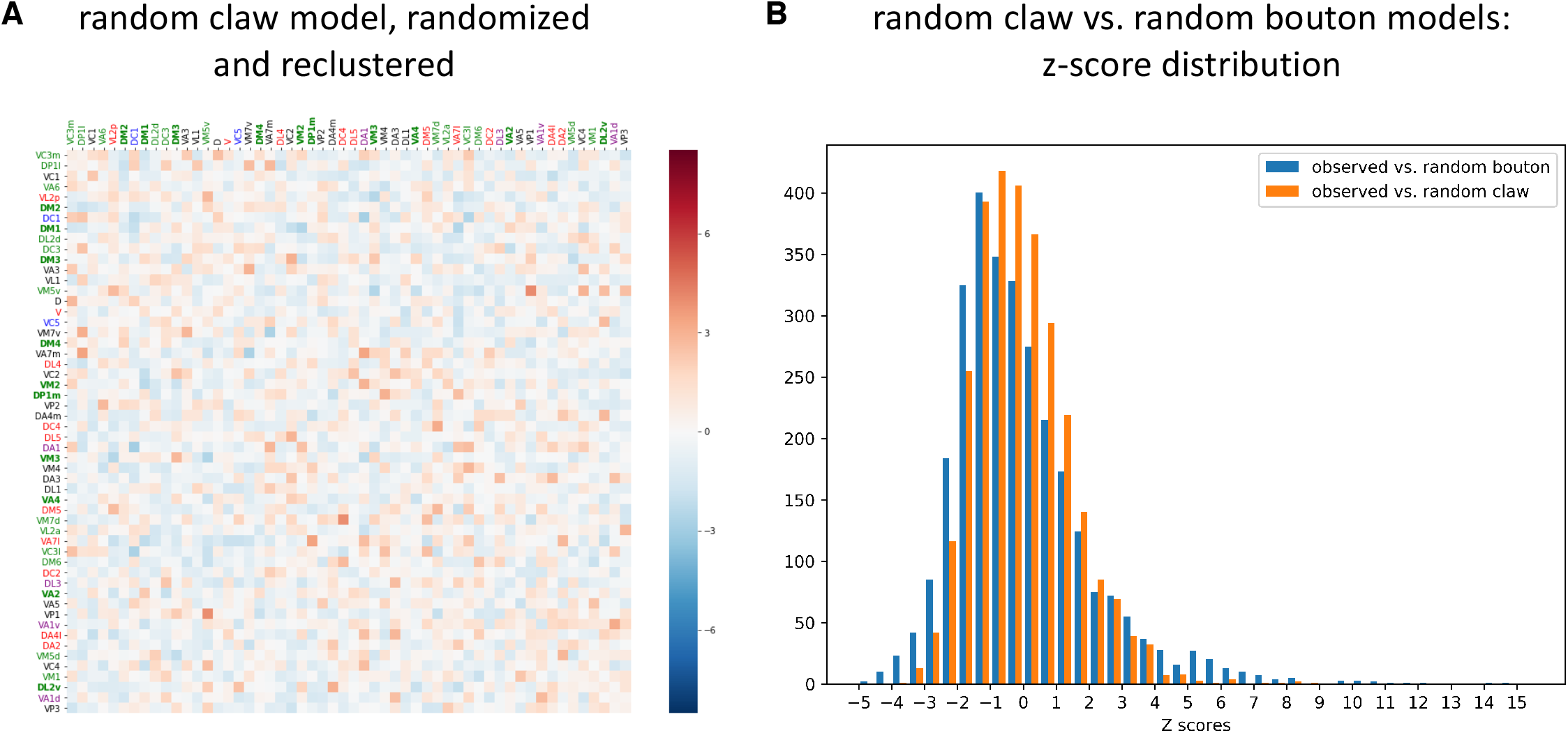
Random claw model. (A) Conditional input analysis was applied to a representative instance of the local random bouton model, with the random bouton model as the null model.Conditional input analysis was applied to a representative instance of the random claw model, with the random claw model as the null model. No cluster of high or low z-scores (2 s.d. or more as compared to the mean of the null model) is seen after K-means re-clustering of the z-score matrix. (B) Distribution of z-scores from conditional input analysis using random bouton null model (z-scores in Figure 1D; i.e. observed vs. random bouton model, mean −0.044, s.d. 2.11) and analysis using the random claw null model (z-scores in Figure 2D; i.e. observed vs. random claw model, mean −0.058, s.d. 1.47). Blue vs. orange distributions, K-S test p < 1×10^-10^. Variance of z-scores is lower using the random claw model than the random bouton model, indicating that the random claw model better captures the observed network structure.

**Supplemental Figure 7.**
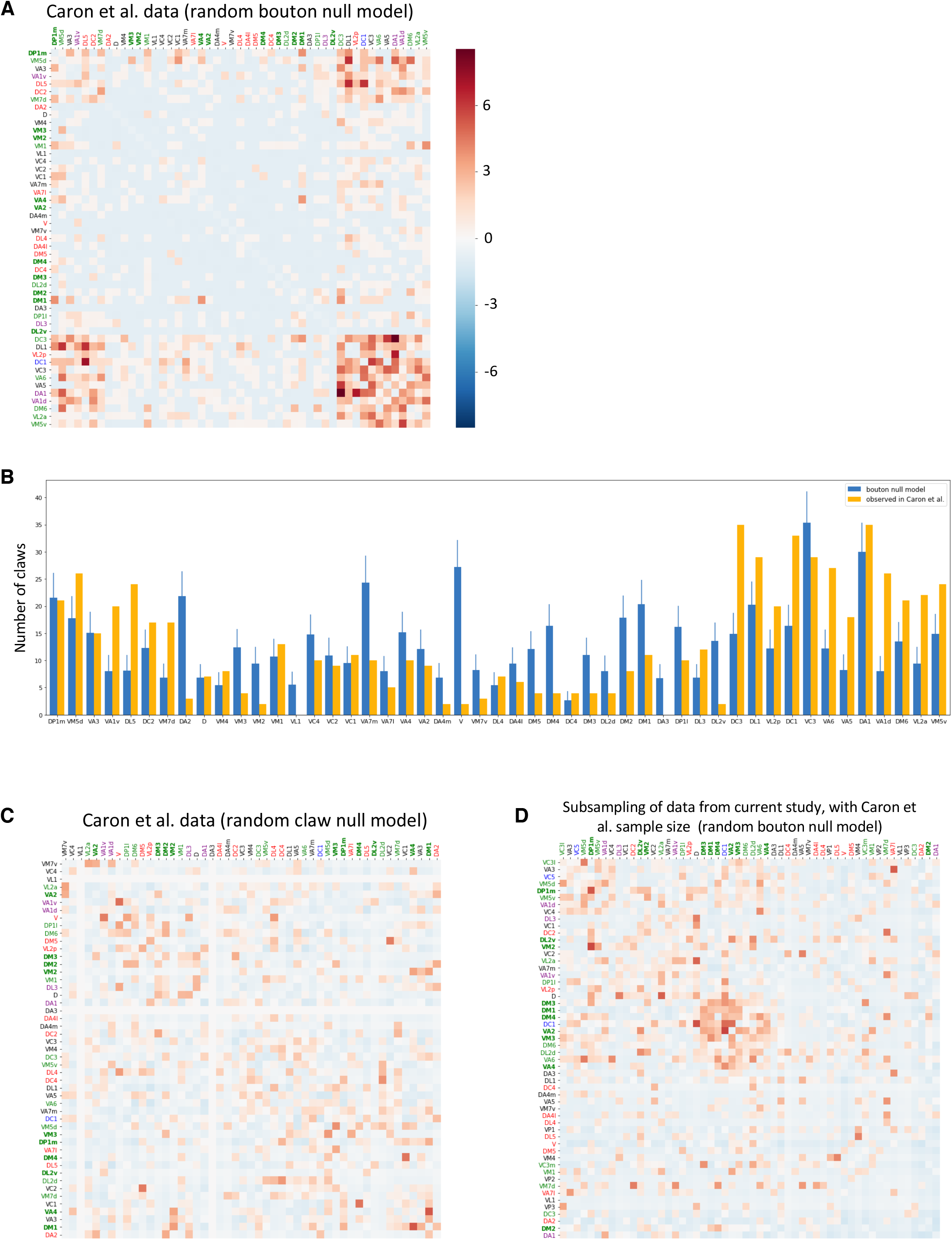
(A) Conditional input analysis of PN-to-KC connectivity data from Caron et al. (2013), using the random bouton null model (Methods). A weak cluster of overconvergent PN types is seen in the lower right corner of the matrix, consistent with the previously reported set of types making the most output onto KCs (Supplemental Figure 1, Caron et al., 2013). This cluster does not overlap strongly with the overconvergent PN community described in the present work (PN type names color coded and bolded as in Figure 1D). (B) Histogram view of data underlying (A). The y-axis shows the mean number of claws receiving input from each PN type in the random bouton null model (blue) and in observed counts the Caron et al. (2013) data (yellow). PN types are ordered as in (A). (C) Conditional input analysis as in (A), except using the random claw null model, which incorporates the observed output rate of each PN type. No overconvergent PN type clusters are discernible using this model. (D) Conditional input analysis of a representatively randomly sampled subset of PN-to-KC connectivity from the current study shows only weak clustering (most z-scores < 2). The number of KCs, and KCs per claw, was held equal to that of the Caron et al. (2013) study.

**Supplemental Figure 8.**
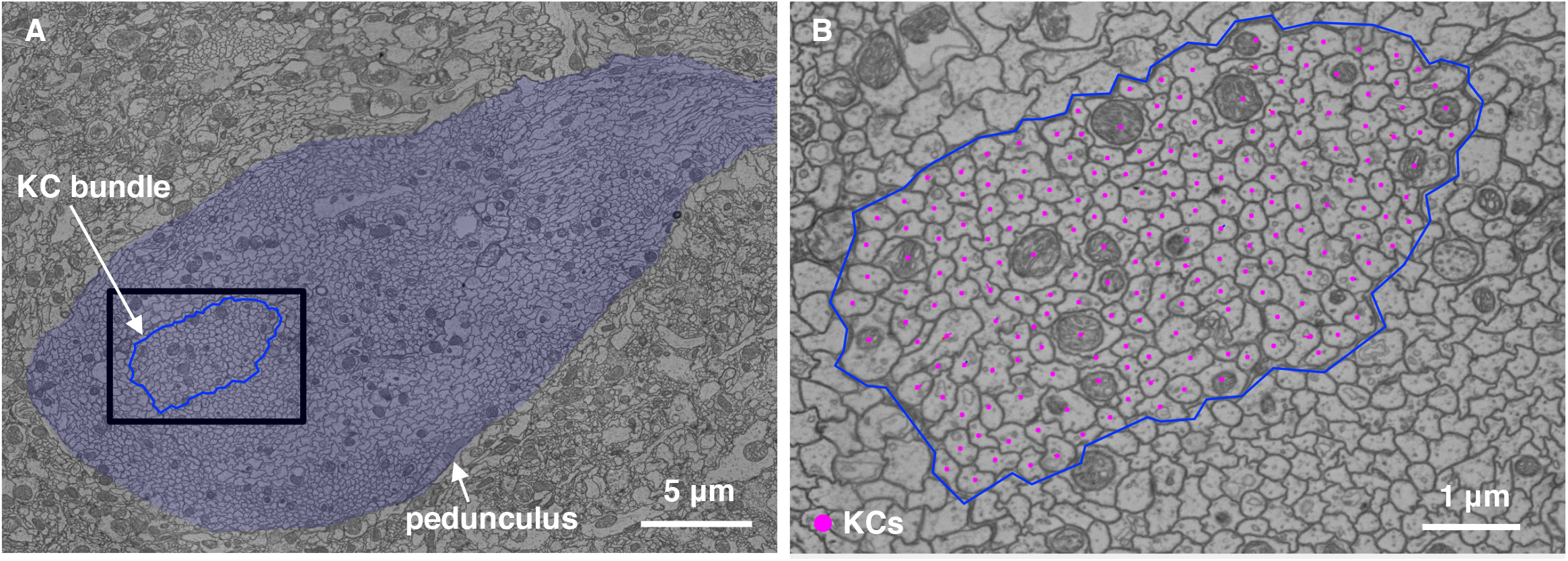
(A) Fasciculating KCs (‘bundle’ KCs) in the pedunculus. A transverse plane image of the pedunculus (shaded blue) shows a discrete bundle of KCs (blue outline) that was completely reconstructed. The black rectangle delineates the subarea shown in (B). (B) Magnified view of the cross-sectional profile of bundle KC axons (magenta dots) in the pedunculus.

**Supplemental Figure 9.**
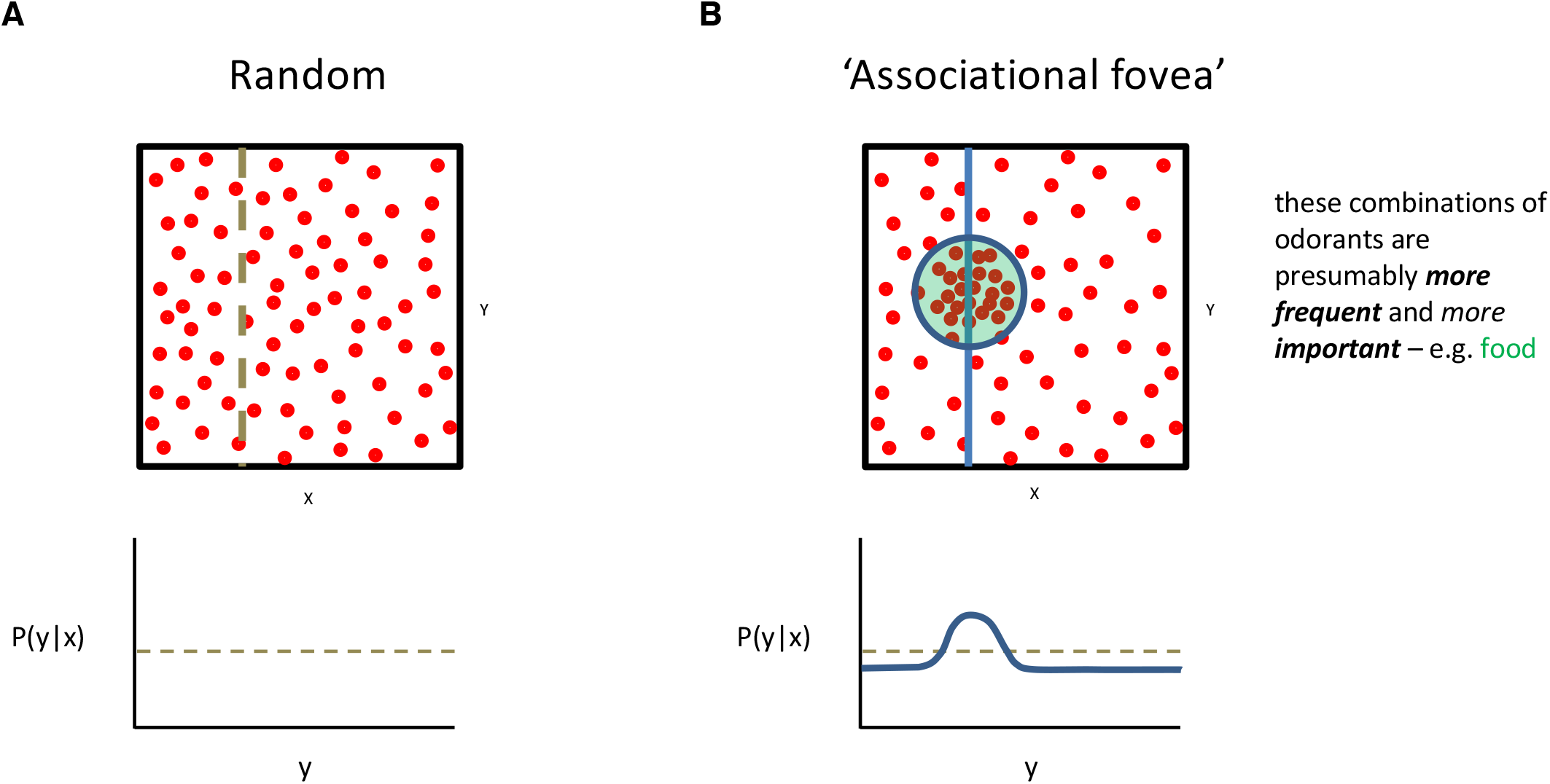
(A) A high dimensional olfactory space is represented schematically here as two-dimensional (x,y axes). Kenyon cells (red dots, upper panel) may be considered as points in this space, with positions defined by their PN inputs. In a random PN-to-KC wiring, the probability that a KC is responsive at a particular position along the y dimension is independent from its responses along the x dimension (lower panel). (B) In the PN-to-KC network structure we observe, KCs receive convergent input from PNs responsive to food-related odorants more often than predicted by chance. Schematically, this may be represented as a non-uniform distribution of KCs within the high dimensional olfactory space defined by PN inputs, analogous to the denser sampling of a visual scene in the fovea of the retina. In this case, the probability that a given KC is responsive to a particular odor along the y axis is *not* independent from whether it is responsive to an odor on the x axis (lower panel). Assuming the fly has a constant number of KCs regardless of network structure (i.e. the number of red dots is the same no matter what), then within the ‘associational fovea’ (green circle, upper panel), the probability is substantially increased, and everywhere else the probability is slightly decreased (lower panel).

**Supplemental Table 1.**

| PN types | Behavioral Significance | Literature |
| --- | --- | --- |
| D | Unknown* | *multiple: aversive (Knaden et al., 2012), pheromonal (Lebreton et al., 2017), and yeast volatiles (ethyl 3-hydroxyhexanoate, Tsakiris et al., 2010) |
| DA1 | Pheromonal | (Kurtovic et al., 2007) |
| DA2 | Aversive | (Stensmyr et al., 2012) |
| DA3 | Unknown |  |
| DA4l | Aversive | (Badel et al., 2016) |
| DA4m | Unknown |  |
| DC1 | Egg-laying | (Dweck et al., 2013) |
| DC2 | Aversive | (Knaden et al., 2012) |
| DC3 | Food | (Ronderos et al., 2014) |
| DC4 | Aversive | (Ai et al., 2010) |
| DL1 | Unknown |  |
| DL2d | Food | (Mansourian and Stensmyr, 2015) |
| DL2v | Food | (Mansourian and Stensmyr, 2015) |
| DL3 | Pheromonal | (van der Goes van Naters and Carlson, 2007) |
| DL4 | Aversive | (Ebrahim et al., 2015) |
| DL5 | Aversive | (Knaden et al., 2012) |
| DM1 | Food | (Sammelhack and Wang, 2009) |
| DM2 | Food | (Schubert et al., 2014) |
| DM3 | Food | (Sammelhack and Wang, 2009) |
| DM4 | Food | (Badel et al., 2016; Semmelhack and Wang, 2009) |
| DM5 | Aversive | (Sammelhack and Wang, 2009) |
| DM6 | Food | (Schubert et al., 2014) |
| DP1l | Food | (Mansourian and Stensmyr, 2015; Silbering et al., 2011) |
| DP1m | Food | (Sammelhack and Wang, 2009) |
| V | Aversive | (Suh et al., 2004) |
| VA1d | Pheromonal | (van der Goes van Naters and Carlson, 2007) |
| VA1v | Pheromonal | (Dweck et al., 2015b) |
| VA2 | Food | (Semmelhack and Wang, 2009) |
| VA3 | Unknown |  |
| VA4 | Food | (Laissue and Vosshall, 2008) |
| VA5 | Unknown |  |
| VA6 | Food | (Mansourian and Stensmyr, 2015; Schlieff and Wilson, 2007) |
| VA7l | Aversive | (Mansourian and Stensmyr, 2015) |
| VA7m | Unknown |  |
| VC1 | Unknown |  |
| VC2 | Unknown¶ | ¶ unclear: dietary antioxidants (Dweck et al., 2015a); suppress oviposition, (Chin et al., 2018) |
| VC3l | Food | (Laissue and Vosshall, 2008) |
| VC3m | Food | (Laissue and Vosshall, 2008) |
| VC4 | Unknown |  |
| VC5 | Egg-laying | (Hussain et al., 2016) |
| VL1 | Unknown |  |
| VL2a | Food | (Grosjean et al., 2011) |
| VL2p | Aversive | (Hamada et al., 2008) |
| VM1 | Food | (Min et al., 2013) |
| VM2 | Food | (Root et al., 2007) |
| VM3 | Food | (Mansourian and Stensmyr, 2015) |
| VM4 | Unknown |  |
| VM5d | Food | (Hallem and Carlson, 2006) |
| VM5v | Food | (Mansourian and Stensmyr, 2015) |
| VM7d | Food | (Laissue and Vosshall, 2008) |
| VM7v | Unknown |  |
| VP1 | Others | temperature (Enjin et al., 2016) |
| VP2 | Others | temperature (Frank et al., 2015; Liu et al., 2015) |
| VP3 | Others | temperature (Enjin et al., 2016) |

## Acknowledgements

We thank: Greg Jefferis and Paavo Huoviala for substantial contributions to the literature search to classify PN types for behavioral significance; Greg Jefferis, Eyal Gruntman, Shaul Druckman, Larry Abbott, Ashok Litwin-Kumar, and Marcus Meister for helpful discussion of preliminary data; Jacob Ratliff, Shahrozia Imtiaz, Benjamin Gorko, Arynne Boyes, Adam John, Emily Moore, Ben Koppenhaver, Philipp Ranft, bailey harrison, Sri Murthy, Ala Haddad, Addy Adesina, Ashley Scott, Chelsea Marlin, Emily Wissell, Zachary Gillis, Saba Ali, Gabrielle Allred, Spencer Waters, Lisa Marin, Annie Scott, Sarah Mohr, Michael Lingelbach, Emma Spillman, Aidan Smith, Teri Ngo, Jordan Dunlap, Bindu Gampah, Melissa Ryan, Nethan Reddy, Adam Fischel, Markus Pleijzier, Arlo Sheridan, Kabas Abou Jahjah, Amelia Edmondson-Stait, Ilenia Salaris, Ruchi Parekh, Austin Warner, Winston Chen, Ruairi Roberts, Julia Gonzales, Laurin Bueld, Cory Ardekani, Razi Rais, Niles Ribeiro, Teresa Neves for Kenyon cell reconstructions; Noah Nelson, for pilot software for analysis of the PN-to-KC connectivity graph at the level of boutons and claws; J. Scott Lauritzen, for help coordinating reconstruction efforts.

## Availability of Source Code and Neuronal Reconstructions

The neuronal reconstructions and source code underlying the analyses presented here are available at: https://github.com/bocklab/pn_kc. Neuronal reconstructions are also available at the Virtual Fly Brain Project (https://v2.virtualflybrain.org/org.geppetto.frontend/geppetto?id=vfb_site/overview.htm).

